# Genome-wide analyses of vocabulary size in infancy and toddlerhood: associations with Attention-Deficit/Hyperactivity Disorder and cognition-related traits

**DOI:** 10.1101/2022.06.01.494306

**Authors:** Ellen Verhoef, Andrea G. Allegrini, Philip R. Jansen, Katherine Lange, Carol A. Wang, Angela T. Morgan, Tarunveer S. Ahluwalia, Christos Symeonides, EAGLE working group, Else Eising, Marie-Christine Franken, Elina Hypponen, Toby Mansell, Mitchell Olislagers, Emina Omerovic, Kaili Rimfeld, Fenja Schlag, Saskia Selzam, Chin Yang Shapland, Henning Tiemeier, Andrew J.O. Whitehouse, Richard Saffery, Klaus Bønnelykke, Sheena Reilly, Craig E. Pennell, Melissa Wake, Charlotte A.M. Cecil, Robert Plomin, Simon E. Fisher, Beate St Pourcain

## Abstract

**Background:** The number of words children produce (expressive vocabulary) and understand (receptive vocabulary) changes rapidly during early development, partially due to genetic factors, although mechanisms are not well understood. Here, we performed a meta-genome-wide association study within the EAGLE consortium and investigated polygenic overlap with later-life traits, including Attention-Deficit/Hyperactivity Disorder (ADHD) and cognition.

**Methods:** We studied 37,913 parent-reported vocabulary size measures (English, Dutch, Danish) for 17,298 children of European descent. Meta-analyses were performed for early-phase expressive (infancy, 15-18 months), late-phase expressive (toddlerhood, 24-38 months) and late-phase receptive (toddlerhood, 24-38 months) vocabulary. Subsequently, we estimated Single-Nucleotide Polymorphism heritability (SNP-h^2^), genetic correlations (r_g_) and modelled underlying genetic factor structures with multivariate models.

**Results:** Contributions of common genetic variation to early-life vocabulary were modest (SNP-h^2^: 0.08(SE=0.01) to 0.24(SE=0.03)) and multi-factorial. Genetic overlap between infant expressive and toddler receptive vocabulary was near zero (r_g_=0.07(SE=0.10)), although both measures were genetically related to toddler expressive vocabulary (r_g_=0.69(SE=0.14) and r_g_=0.67(SE=0.16), respectively). Consistently, polygenic association patterns with later-life traits differed: Genetic links with cognition emerged only in toddlerhood (e.g. toddler receptive vocabulary and intelligence: r_g_=0.36(SE=0.12)), despite comparable study power for infant measures. Furthermore, increased polygenic ADHD risk was associated with larger infant expressive vocabulary (r_g_=0.23(SE=0.08)), as confirmed by ADHD-symptom-based follow-up analyses in the Avon Longitudinal Study of Parents and Children (ALSPAC-r_g_=0.54(SE=0.26)). Genetic relationships with toddler receptive vocabulary were, however, opposite (ALSPAC-r_g_=-0.74(SE=0.23)), highlighting developmental changes in genetic architectures.

**Conclusions:** Multiple genetic components contribute to early-life vocabulary development, shaping polygenic association patterns with later-life ADHD symptoms and cognition.

## Introduction

Language development in infants and toddlers is often assessed with measures of expressive and receptive vocabulary^1,2^. These constructs relate to language production and understanding, respectively, and can be measured relatively easily (albeit indirectly) through parental reports. The first spoken words, representing one of the milestones in language development, typically emerge between the ages of 10 to 15 months^2^. Receptive vocabulary development usually precedes expressive vocabulary development, emerging at six to nine months of age^3^. Consequently, the number of words children understand is often larger than the number of words they produce, and exceeds the latter at least four-fold based on parent-reported measures at 16 months of age^4^. Once children reach an expressive vocabulary size of ~50 words at the age of 12 to 18 months, there is often a period of rapid vocabulary growth around 16 to 22 months of age^5^, resulting in a vocabulary size of 100 to 600 words at 24 months^4^.

During the early stages of language learning (infancy, ≤18 months of age) children typically produce words in isolation^2^, followed by a period of acquiring two-word combinations and more complex grammatical structures^4,6^. Across stages, there are moderate-to-strong phenotypic correlations between measures of vocabulary size assessed at one-year intervals (r_p_=0.47-0.63)^7,8^. Such correlations tend, however, to decrease with increasing age windows^8^, suggesting phenotypic heterogeneity across development. In addition, phenotypic correlations between different language constructs assessed at the same age often exceed phenotypic correlations for the same construct at different ages^7^.

Individual differences in early-life vocabulary development can, partially, be explained by genetic factors^7,9–11^. Twin heritability estimates for expressive vocabulary range between 10% and 25% (24-36 months)^7,9,10^, reflecting phenotypic variation due to all possible genetic influences. These findings are corroborated by genetic research investigating Single-Nucleotide Polymorphisms (SNPs), with SNP-h^2^ estimates of 13% and 14% (15-30 months)^9^. For receptive vocabulary, a twin heritability estimate of 28% was reported at 14 months of age^12^. Evidence for SNP-h^2^ at a similar age was poor but was present at 38 months of age (SNP-h^2^=12%)^8^.

Population-based studies of English-speaking children showed that the genetic architecture of language development spanning infancy to early childhood is complex, with evidence for both stability and change in underlying genetic contributions^7–9,13^. At the genome-wide level, genetic correlations (r_g_) for expressive vocabulary measures between 15 and 38 months of age ranged from 0.48 to 0.74^7–9^, suggesting only moderate-to-strong genetic stability. At the individual SNP level, a previous meta-genome-wide association study (meta-GWAS, N=8,889) identified rs7642482 on chromosome 3p12.3, near the *ROBO2* gene^9^, to be associated with expressive vocabulary in infants (age: 15-18 months) but attenuated in toddlers (age: 24-30 months)^9^. These changes might be due to age-specific genetic mechanisms, potentially reflecting phenotypic heterogeneity, highlighting the need for more powerful studies to identify and characterise genetic associations.

Genetic influences underlying early-life vocabulary are shared with many later childhood abilities. In UK twins, for example, early expressive language skills (24-48 months) were moderately genetically correlated (r_g_=0.36) with childhood reading abilities^13^. Similarly, in a UK population-based genomic study, receptive vocabulary size (38 months) showed moderate-to-strong genetic correlations (r_g_=0.58-0.92) with mid-childhood reading skills^14^. The genetic link of infant and toddler vocabulary size with cognition-related skills assessed beyond mid-childhood is, however, not fully understood and may reveal an early manifestation of later-life cognitive functioning.

Genetic influences underlying early-life vocabulary may also be shared with other behavioural and health measures, including childhood-onset neurodevelopmental conditions such as Attention-Deficit/Hyperactivity Disorder (ADHD) and Autism Spectrum Disorder (ASD). For example, children with ADHD often experience difficulties with mastering language and literacy skills^15–17^. Furthermore, poor language skills at the age of three years were found to be predictive of inattention and hyperactive symptoms two years later in life^18^. More specifically, there is evidence for genetic overlap between higher ADHD risk and lower mid-childhood/early-adolescent language- and literacy-related abilities, primarily implicating reading performance^19–22^. For children diagnosed with ASD, the phenotypic spectrum is broader, including children with little or no spontaneous spoken language by the time they reach school age^23^, as well as individuals with, comparably, few problems in the language domain^24^. Despite the phenotypic overlap between neurodevelopmental disorder and language traits, little is known about genetic links across different stages during language development. In particular, establishing genetic relationships with developmental language symptoms preceding the typical age of onset for neurodevelopmental conditions in toddlerhood or childhood^25,26^ may, eventually, contribute to an earlier identification of disorder-related symptoms. In addition, genetic links between children’s vocabulary and head circumference measures may inform on abnormal brain development^27^, given strong correlations with MRI brain volume^28,29^.

In this study, we aim to elucidate the polygenic architecture of vocabulary acquisition. We investigate developmental changes in genetic contributions to vocabulary at the single-variant and trait covariance level by performing meta-GWASs of expressive and receptive vocabulary size at different developmental stages: Studied developmental windows include an early, single-word phase (15-18 months, infancy) and a late phase during which children start using two-word combinations and more complex grammatical structures (24-38 months, toddlerhood). Fitting bi-variate and multi-variate structural models, we report polygenic links with childhood behavioural and health measures, as well as adult cognition-related outcomes.

## Methods and Materials

### Phenotype selection and study design

Cohorts with quantitative vocabulary scores assessed during the first three years of life and genome-wide genotypes were invited to participate in this study, embedded within the EArly Genetics and Life Course Epidemiology (EAGLE) consortium^30^ (https://www.eagle-consortium.org/working-groups/behaviour-and-cognition/early-language/). Expressive vocabulary scores were assessed between 15 and 38 months of age and analysed across two developmental stages to allow for age-specific genetic influences: an early phase (15-18 months, infancy) and a late phase (24-38 months, toddlerhood). Scores for receptive vocabulary were included for the late phase only due to limited data availability, low reliability and little evidence for SNP-h^2^ during the early phase^8^ (Supplemental Methods).

Up to seven population-based cohorts (Supplemental Methods) participated in this study, of which two had longitudinal vocabulary assessments (Figure 1, Table S1). Vocabulary scores were ascertained by parental report using age-specific word lists that were adapted from the MacArthur Communicative Development Inventory (CDI)^10,31–35^ or the Language Development Survey (LDS)^36^ (Supplemental Methods, Table S1). Ethical approval was obtained by the local research ethics committee for each participating study, and all parents and/or legal guardians provided written informed consent (Supplemental Methods).

**Figure 1:**
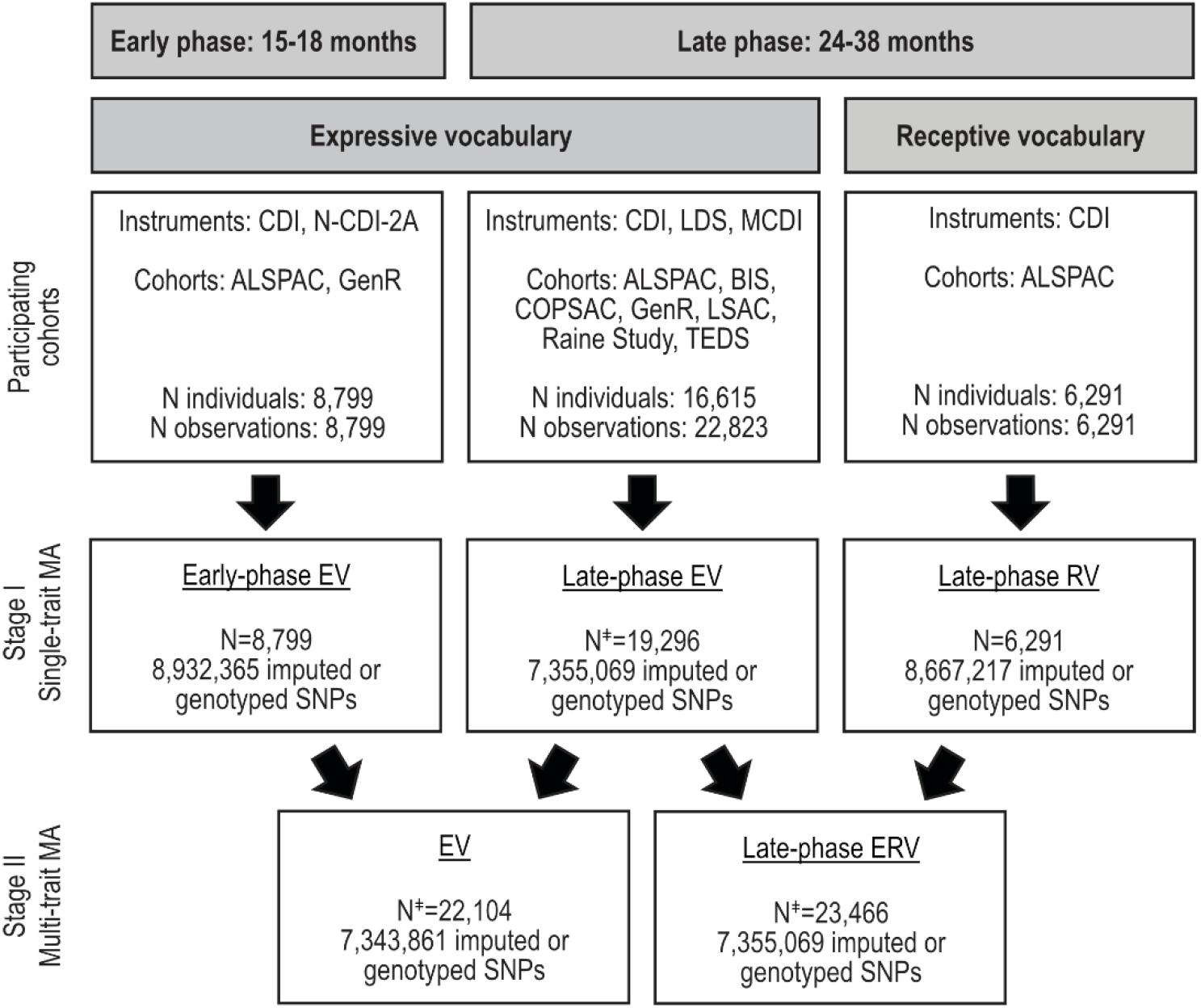
Meta-analysis study design. Vocabulary size was assessed between 15-38 months of age and studied with respect to an early (15-18 months, infancy) and late phase (24-38 months, toddlerhood) of language acquisition to allow for age-specific genetic influences. Scores for receptive vocabulary were included in the late-phase only. In stage I, three single-trait meta-analyses were conducted: early-phase expressive vocabulary, late-phase expressive vocabulary and late-phase receptive vocabulary. In stage II, multi-trait genome-wide analyses were performed across early-phase and late-phase expressive vocabulary, as well as across late-phase expressive and receptive vocabulary to increase statistical power. ‡ Estimated sample size based on the increase in mean χ^2^ statistic using multi-trait analysis of genome-wide association. Abbreviations: ALSPAC, Avon Longitudinal Study of Parents and Children; BIS, Barwon Infant Study; CDI, Communicative Development Inventory; COPSAC, Copenhagen Prospective Studies on Asthma in Childhood; EV, expressive vocabulary; ERV, expressive and receptive vocabulary; GenR, Generation R Study; LDS; Language Development Survey; LSAC, Longitudinal Study of Australian Children; MA, meta-analysis; RV, receptive vocabulary; TEDS, Twins Early Development Study

### Genotyping and imputation

Genotyping was conducted using high-density SNP arrays within each cohort, and quality control followed standard procedures^37^ (Table S2). In total, between 440,476 and 608,517 high-quality autosomal genotyped markers were imputed against the Haplotype Reference Consortium (HRC) r1.1 panel^38^ (Table S2).

### Single variant association analyses and meta-analyses

Within each cohort, vocabulary scores were adjusted for potential covariates (Supplemental Methods), and rank-transformed to achieve normality and to allow for comparisons of genetic association effects across different psychological instruments. SNP-vocabulary associations were then estimated within each cohort using linear regression of rank-transformed residuals on posterior genotype probability, assuming an additive genetic model, except for the Longitudinal Study of Australian Children (LSAC). Within LSAC, best-guess genotypes were analysed assuming an additive genetic model (Supplemental Methods). Prior to meta-analysis, GWAS summary statistics underwent extensive quality control using the EasyQC R package^39^ (v9.2) (Table S2, Supplemental Methods).

As part of analysis stage I (Figure 1), single-trait meta-analyses were performed for early-phase expressive vocabulary, late-phase expressive vocabulary and late-phase receptive vocabulary using either METAL^40^ and/or multi-trait analysis of genome-wide association (MTAG)^41^ software (Supplemental Methods). As part of analysis stage II (Figure 1), multi-trait meta-analyses were performed with MTAG^41^, combining genetically correlated vocabulary summary statistics to increase statistical power.

The number of independent vocabulary measures analysed in this study was estimated with a Matrix Spectral Decomposition (matSpD)^42^ based on bivariate genetic correlations (see below) across the three single-trait vocabulary meta-analyses (stage I). Adjusting the genome-wide association significance threshold for 2.38 independent vocabulary measures resulted in a multiple-testing-adjusted genome-wide association significance threshold of *P*<2.10×10^-8^ (5×10^-8^/2.38).

### FUMA analyses

SNP-vocabulary associations passing the unadjusted genome-wide significance threshold (*P*<5×10^-8^) were identified and annotated using Functional Mapping and Annotation of genetic associations^43^ software (FUMA v1.3.6). In addition, gene-based genome-wide, gene-set and gene-property analyses were conducted with Multi-marker Analysis of GenoMic Annotation (MAGMA, v1.08) within FUMA^43^ (v1.3.6a) (Supplemental Methods).

### SNP-heritability and genetic relationship analyses

SNP-h^2^ was estimated for all vocabulary GWAS summary statistics created in this study (stage I and II) and for traits that were screened for bivariate genetic correlation (r_g_) with vocabulary size (see below), using High-Definition Likelihood^44^ (HDL, Supplemental Methods). HDL software estimates SNP-h^2^ and r_g_ with increased accuracy compared to Linkage Disequilibrium Score (LDSC) regression analyses^44^. To confirm the robustness of HDL results, SNP-h^2^ was also estimated using LDSC regression^45^ for vocabulary summary statistics.

To ensure sufficient power for HDL-r_g_ analyses, we selected traits with evidence for HDL-SNP-h^2^ (*P*<0.05) and HDL-SNP-h^2^ Z-score >4^46^ (Supplemental Methods). We assessed genetic overlap (i) among single-trait vocabulary measures (stage I) and (ii) across single-trait vocabulary measures (stage I) and several health-, cognition-, and behaviour-related outcomes: intelligence^47^ (5-98 years, N=279,930), educational attainment^48^ (>30 years, N=766,345), infant head circumference^49^ (6-30 months, N=10,768), childhood head circumference^50^ (6-9 years, N=10,600), childhood aggressive behaviour^51^ (1.5-18 years, N=151,741), childhood internalising symptoms^52^ (3-18 years, N=64,641), ADHD^53^ (N=53,293; N_cases_=19,099) and ASD^54^ (N=46,350; N_cases_=18,381). The multiple-testing-adjusted threshold for HDL-r_g_ analyses was defined at 6.59×10^-3^. This threshold reflects a correction for 7.59 independent traits considered (0.05/7.59), as estimated using matSpD^42,55^ and a bivariate genetic correlation matrix (Figure S1).

Polygenic prediction of late-phase expressive vocabulary was carried out using polygenic scoring (Supplemental Methods).

### Structural equation modelling

To obtain insight into the developmentally changing genetic covariance patterns of ADHD with vocabulary size during infancy and toddlerhood, we modelled the underlying multivariate genetic architecture. We selected an individual-level data approach for this, as summary statistic-based SEM modelling using Genomic SEM^56^ was not feasible due to a violation of modelling assumptions (see Supplementary Methods). Individual-level data from unrelated children of the Avon Longitudinal Study of Parents And Children (ALSPAC) cohort^57,58^ (N≤6,524) was studied using genetic-relationship-matrix structural equation modelling^59^ (grmsem, v1.1.2) (Supplemental Methods). ALSPAC measures of vocabulary size were identical to those included in the meta-GWAS (Table S1). ADHD symptom expression scores were assessed with the Strengths and Difficulties Questionnaire^60^, based on teacher and mother reports between 7 and 17 years of age. For both teacher- and mother-report, the ADHD symptom score with the highest SNP-h^2^ Z-score was selected for subsequent analyses (Supplementary Information, Table S3). Given the temporal order of studied traits, we fitted a GRM-SEM Cholesky decomposition^61^ to the data. A Cholesky decomposition dissects the phenotypic covariance structure into additive genetic factors (A), capturing genetic variance tagged by common genotyped SNPs^59^, and residual factors (E), reflecting all other sources of variance, including error. It is a saturated model with as many latent genetic and residual factors as observed variables, without any restrictions on the structure^61^. Subsequently, genetic and residual correlations were estimated according to theory^62^ using grmsem. Phenotypic correlations (r_p_) were derived using Pearson correlation in R (R:stats library, v.4.1.0).

## Results

### Single-trait and multi-trait meta-GWAS

Single-trait genome-wide association analyses were carried out for early-phase expressive (15-18 months, N=8,799), late-phase expressive (24-38 months, N=16,615) and late-phase receptive (24-38 months, N=6,291) vocabulary size (stage I, Figure 1) using data from English-, Dutch- or Danish-speaking children of European descent from seven independent cohorts (Table S1). There was little evidence for novel SNP signals at the multiple-testing-adjusted genome-wide significance level (*P*<2.10×10^-8^, Figure S2a-c). For early-phase expressive vocabulary, a single GWAS signal passed the unadjusted genome-wide significance threshold (rs9854781, *P*<5×10^-8^), consistent with a known locus identified through a previous meta-GWAS studying overlapping samples (rs764282, LD-r^2^=0.78)^9^. Genome-wide gene-based, gene-set and gene-property analyses with MAGMA^63^ did not provide evidence for association passing the multiple-testing-adjusted significance thresholds (Figure S3, Table S4).

All early-life vocabulary measures were modestly heritable, with SNP-h^2^ estimates of 0.24(SE=0.02), 0.08(SE=0.01), and 0.20(SE=0.04) for early-phase expressive vocabulary, late-phase expressive vocabulary, and late-phase receptive vocabulary, respectively (Figure 2a, Table S5). Given limited data availability, polygenic prediction (out of the meta-analysis samples) was carried out for late-phase expressive vocabulary only (β=0.04(SE=0.04), *P*=0.35, R^2^=0.14%). Power to detect genetic overlap was, however, low (≤0.11) due to a combination of low SNP-h^2^ and low target sample size (Early Language in Victoria Study^64^, N=639).

**Figure 2:**
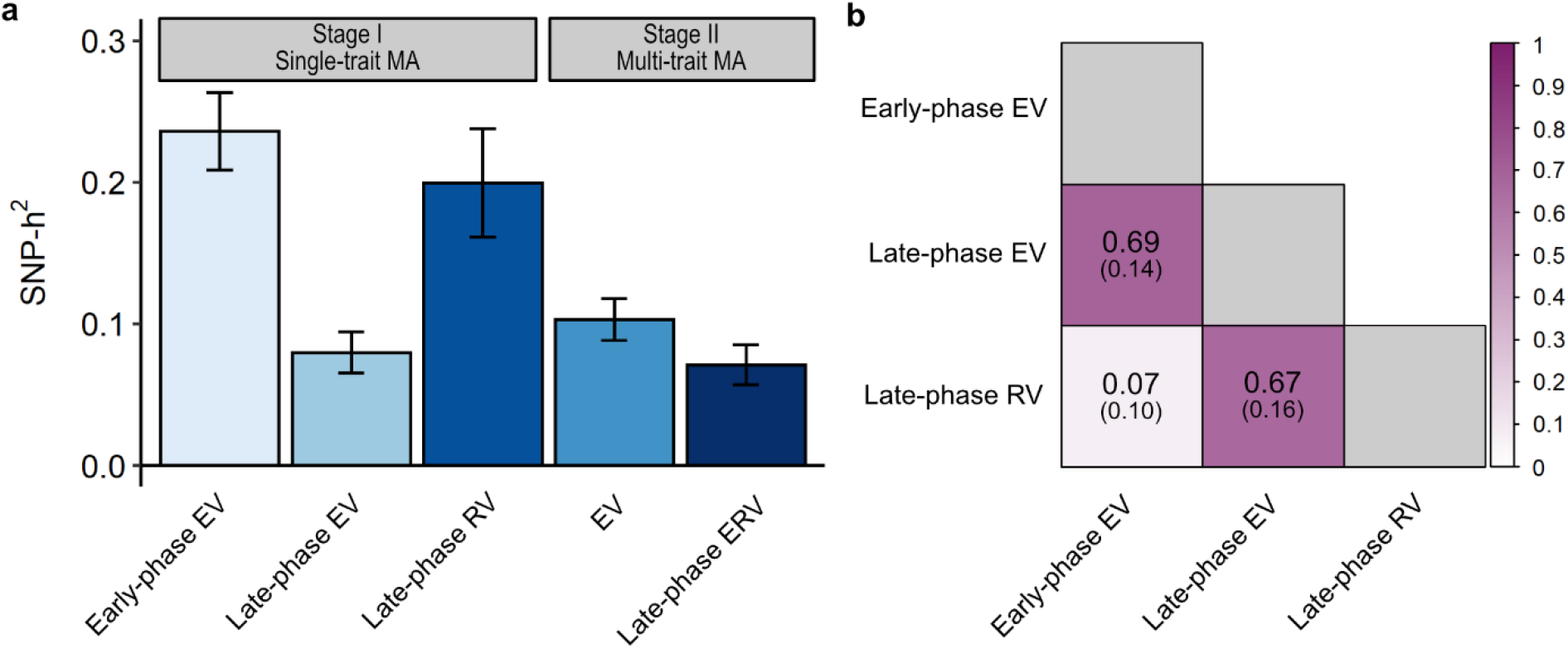
SNP-heritability and genetic correlations of vocabulary size. **(a)** SNP-heritability estimates for single- and multi-trait vocabulary summary statistics were estimated with High-Definition Likelihood^44^ software. Error bars represent standard errors. **(b)** Genetic correlations (r_g_) among single-trait vocabulary summary statistics were estimated with High-Definition Likelihood^44^ software. Corresponding standard errors are shown in brackets. Abbreviations: EV, expressive vocabulary; ERV; expressive and receptive vocabulary; MA, meta-analyses; RV, receptive vocabulary; SNP-h^2^, Single-Nucleotide Polymorphism heritability

Genetic correlations between early- and late-phase expressive vocabulary (r_g_=0.69(SE=0.14)), as well as between late-phase expressive and receptive vocabulary (r_g_=0.67(SE=0.16)), were moderate (Figure 2b), suggesting some stability in genetic factors during development. Genetic influences underlying early-phase expressive vocabulary were, however, largely independent of those related to late-phase receptive vocabulary (r_g_=0.07(SE=0.10)). Given comparable statistical power to detect genetic overlap with late-phase receptive vocabulary (e.g. statistical power at r_g_=0.70: early-phase expressive vocabulary=83%, late-phase expressive vocabulary=71%), these findings suggest developmental genetic heterogeneity and the existence of at least two independent genetic factors.

To maximise statistical power for single-variant discovery, genetically correlated vocabulary measures were combined as part of two multi-trait meta-analyses using MTAG (stage II, Figure 1). However, we neither identified further SNP-vocabulary associations (Figure S2d-e, Table S6) nor increased evidence for SNP-h^2^ (Z-scores, Figure 2a, Table S5), and subsequent analyses were restricted to stage I vocabulary summary statistics only.

### Genetic relationships with health-, cognition- and behaviour-related outcomes

We investigated genetic links between early-life vocabulary measures (stage I) and several heritable health-, cognition- and behaviour-related outcomes (see Table S7 for SNP-h^2^) by estimating genetic correlations with HDL software^44^ (multiple-testing-adjusted threshold: *P*<6.59×10^-3^). Consistent with the estimated genetic architecture underlying early-life vocabulary size (see above), polygenic association patterns for early-phase and late-phase vocabulary were highly divergent (Figure 3a). While there was little evidence for genetic links with cognition during infancy, positive genetic correlations between expressive and receptive vocabulary emerged only during toddlerhood (Figure 3a), despite comparable analysis power across both developmental stages (Table S8). Both, larger late-phase expressive and receptive vocabulary size were genetically correlated with higher intelligence across the lifespan (late-phase expressive vocabulary: r_g_=0.32(SE=0.08); late-phase receptive vocabulary: r_g_=0.36(SE=0.12)) and higher adult educational attainment (late-phase expressive vocabulary: r_g_=0.26(SE=0.05); late-phase receptive vocabulary: r_g_=0.37(SE=0.06)). In addition, genetic relationships with childhood behaviour-related traits, such as ADHD, changed during development (Figure 3a). While larger early-phase expressive vocabulary size was genetically correlated with increased ADHD risk (r_g_=0.23(SE=0.08)), and, at the nominal level (*P*<0.05), childhood aggressive behaviour (r_g_=0.42(SE=0.16)), such genetic associations were attenuated for both late-phase expressive and receptive vocabulary measures (Figure 3a).

**Figure 3:**
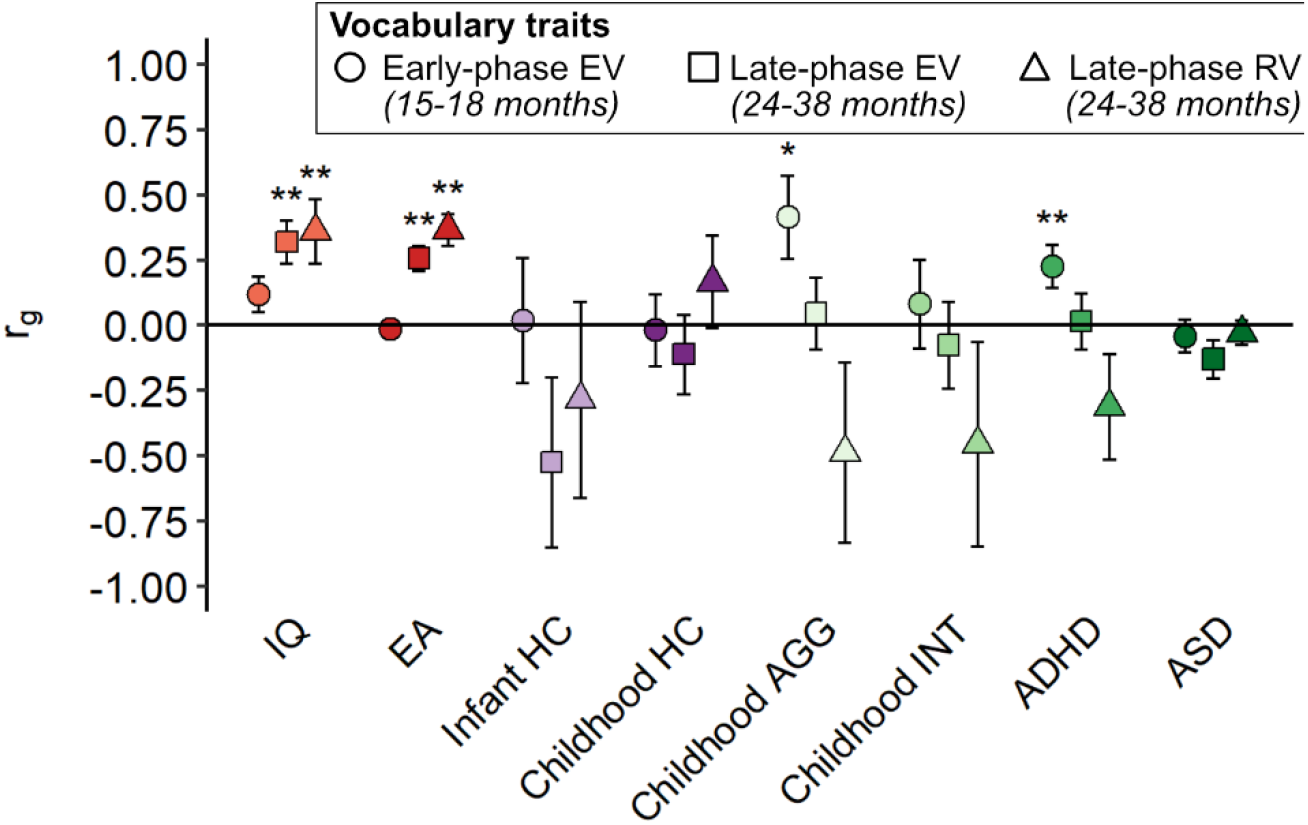
Genetic correlations of vocabulary with several health, cognitive, and behavioural outcomes. Genetic correlations (r_g_) were estimated using summary statistics and High-Definition Likelihood (HDL)^44^. Bars represent standard errors. ** multiple-testing adjusted *P*<6.59×10^-3^; * *P*<0.05. Abbreviations: ADHD, Attention-Deficit/Hyperactivity Disorder; AGG, aggression; AIC, Akaike Information Criterium; ASD, Autism Spectrum Disorder; CFI, comparative fit index; df, degrees of freedom; EA, educational attainment; EV, expressive vocabulary; HC, head circumference; INT, internalising symptoms; IQ, general intelligence; M, months; RV, receptive vocabulary; SRMR, standardised root mean squared residual

To explore developmental changes in genetic association patterns with ADHD symptoms *in depth,* we studied individual-level data from children of the ALSPAC cohort using GRM-SEM^59^. Note that it was not possible to fit an informative summary statistic-based SEM model with HDL-based genomic SEM^56^, as the studied summary statistics did not fulfil modelling requirements (see Supplementary Methods).

Specifically, we dissected the phenotypic covariance of early-life vocabulary measures and ADHD symptoms at ages 8 and 13 years (selected based on the highest SNP-h^2^ Z-score, Table S3) into genetic (A) and residual (E) factors by fitting a saturated (Cholesky) structural model (Figure 4a, Table S9). Subsequently, we studied Cholesky-model-derived genetic correlations (Figure 4b). Confirming HDL findings, larger early-phase expressive vocabulary (15 months) was genetically correlated with more ADHD symptoms (r_g_ADHD8y_=0.56(SE=0.26); r_g_ADHD13y_=0.54(SE=0.25), Figure 4b). This association was captured by a genetic factor A1, with positive factor loadings (λ) for early-phase expressive vocabulary and ADHD symptoms at 8 and 13 years (λ_exp_voc15m_=0.34(SE=0.07); λ_ADHD8y_=0.28(SE=0.13); λ_ADHD13y_=0.27(SE=0.12)). In contrast, we observed an inverse genetic correlation between larger late-phase receptive vocabulary size (38 months) and lower ADHD symptoms (r_g_ADHD8y_=-0.60(SE=0.23); r_g_ADHD13y_=-0.74(SE=0.16), Figure 4b). This inverse genetic association was captured by two independent genetic factors, A2 and A4 (Figure 4a). A2 reflects genetic influences underlying expressive vocabulary size at 24 months (λ_exp_voc24m_=0.33(SE=0.06)), independent of A1, that were inversely linked to ADHD symptoms at 8 and 13 years (λ_ADHD8y_=-0.41(SE=0.11); λ_ADHD13y_=-0.25(SE=0.12)). A4 explained unique genetic variance contributing to receptive vocabulary size at 38 months (λ_rec_voc38m_=0.15(SE=0.08), although not passing the conventional level of significance with *P*=0.07) and was inversely associated with ADHD symptoms at 13 years (λ_ADHD13y_=-0.34(SE=0.14)). Thus, the Cholesky-estimated genetic factor structures revealed opposite ADHD association patterns for early-phase (infancy) versus late-phase (toddlerhood) vocabulary measures (Figure 4a, Table S9).

**Figure 4:**
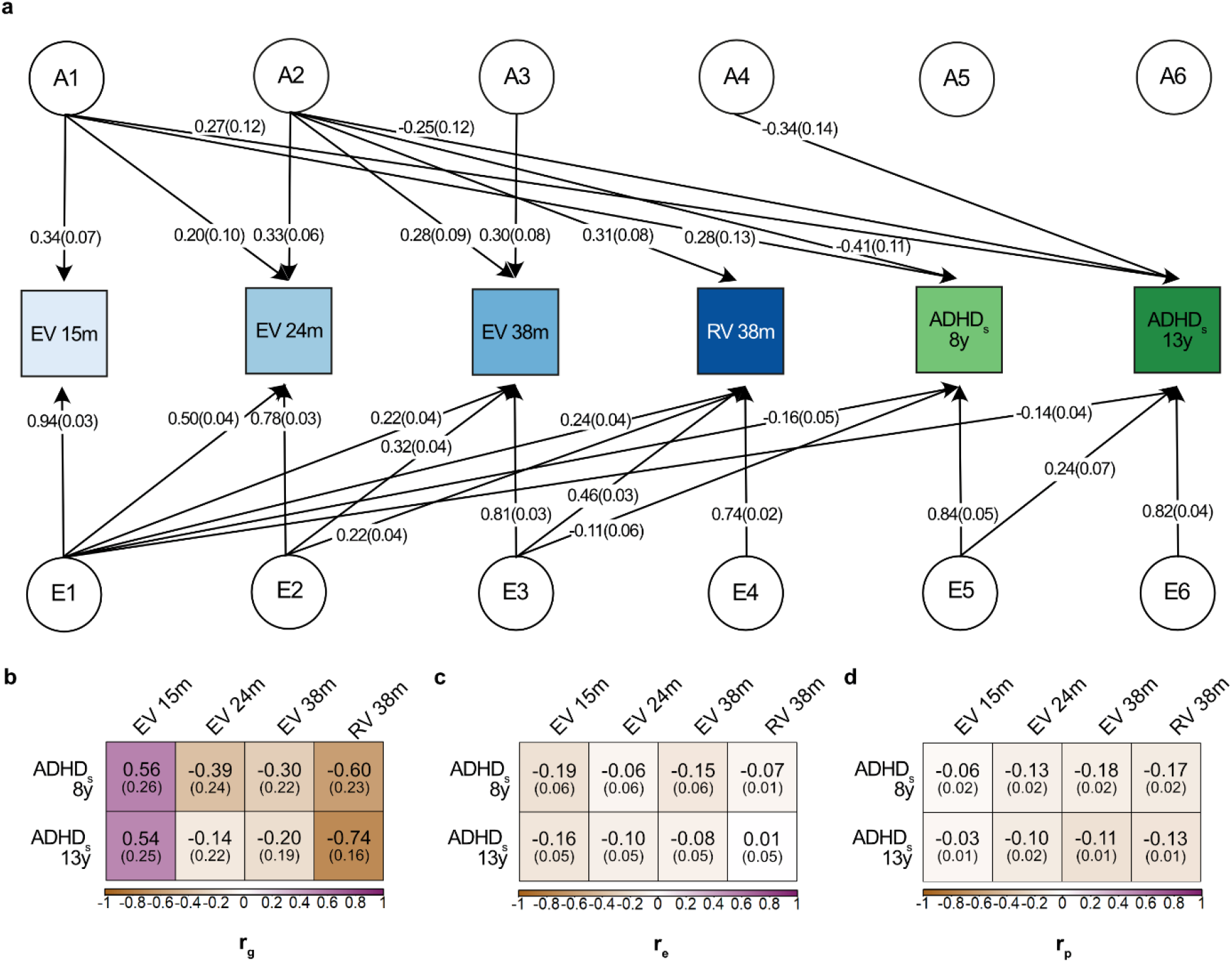
Cholesky decomposition of early-life vocabulary size and later ADHD symptoms. Genetic-relationship matrix structural equation modelling (GRM-SEM) of vocabulary scores (15, 24 and 38 months of age) in combination with ADHD symptom expression scores (teacher-report at 8 years and mother-report at 13 years) based on all available observations for children across development (N≤6,524). Individual-level data was retrieved from the Avon Longitudinal Study of Parents and Children. **(a)** Path diagram with standardised path coefficients and corresponding standard errors for a Cholesky decomposition. Only paths with a path coefficient of *P*<0.05 are shown. Full information on all path coefficients and their standard errors can be found in Table S9. **(b)** Genetic, **(c)** residual and **(d)** phenotypic correlation patterns between early-life vocabulary size and ADHD symptom scores assessed at 8 and 13 years of age. Genetic (r_g_) and residual (r_e_) correlations were estimated with genetic-relationship-matrix structural equation modelling^59^ based on a Cholesky decomposition (shown in a). Phenotypic correlations (r_p_) were estimated with Pearson correlation. Standard errors are shown in parentheses. Abbreviations: ADHDs, Attention-Deficit/Hyperactivity Disorder symptoms; EV, expressive vocabulary; m, months; r_e_, residual correlation; r_g_, genetic correlation; r_p_, phenotypic correlation; RV, receptive vocabulary; SDQ, Strengths and Difficulties Questionnaire; y, years

Next, we compared phenotypic (r_p_) and Cholesky-derived residual (r_e_) correlations in the ALSPAC sample. While genetic, residual and phenotypic correlations had the same direction of effect for all three late-phase vocabulary measures (Figure 4b-d), we uncovered a rare violation of Cheverud’s conjecture^65^. in infancy. Cheverud’s conjecture postulates that phenotypic correlations are likely to be fair estimates of their genetic counterparts^65^. In infancy, however, we observed a positive genetic association between early-phase expressive vocabulary and ADHD symptoms (r_g_ADHD8y_=0.56(SE=0.26)) that was masked at the phenotypic level (r_p_ADHD8y_=-0.06(SE=0.02)) by a negative residual correlation (r_e_ADHD8y_=-0.19(SE=0.06)), as shown here for teacher-reported ADHD scores at age 8 years.

## Discussion

This meta-GWAS of expressive and receptive vocabulary size in infancy and toddlerhood identified marked differences in genetic influences underlying vocabulary measures at different developmental stages, as demonstrated with genomic covariance modelling. The heterogeneity within the genetic architecture underlying early-life vocabulary size matched distinct polygenic association patterns with ADHD and cognition-related traits.

Bivariate genetic correlation patterns and multivariate structural models suggested at least two independent genetic factors contributing to early-life vocabulary size, confirming previous reports of a heterogeneous genetic architecture^7–9,13^. Findings are also consistent with developmentally increasing phenotypic heterogeneity across widening age windows^8^. Genetic influences contributing to utterances in infancy, approximated here by early-phase expressive vocabulary size (15-18 months), may capture the first stages of language learning. During this phase of “learning to speak”, speech is emerging, and words are usually produced in isolation^2^. More specifically, children acquire phonological skills to identify phonemes and sequences from speech and store them for future production^66^, but also develop oral motor^67^ and speech motor skills^68^. Despite sufficient statistical power (with >90% power to detect r_g_ ≥0.20 with EA), there was little evidence for genetic overlap between early-phase vocabulary scores and adult EA and cognition. Such associations only emerged for late-phase vocabulary scores during toddlerhood (24-38 months), in line with previous work studying mid-childhood cognition-related measures^8^, and suggest specificity. The genetic overlap of toddler vocabulary measures with cognition may reflect the onset of a subsequent phase of “speaking to learn”, where toddlers have mastered some language fluency and started to use word combinations and more complex grammatical structures^4,6^. Together, our findings highlight rapid changes in the genetic architecture of vocabulary acquisition across a period of less than two years, even when assessed with similar psychological instruments. Shared genetic influences across both developmental phases, illustrated by the strong genetic overlap of late-phase expressive vocabulary with both early-phase expressive and late-phase receptive vocabulary, underline the dynamic character of this process.

Heterogeneity in genetic components contributing to early-life vocabulary size was further reflected by distinct polygenic association patterns with later-life behavioural traits. During infancy, larger expressive vocabulary was associated with an increased polygenic risk for ADHD. Consistently, younger age at first walking^69^, another early developmental milestone, and better infant gross motor skills^70^ have been linked to higher polygenic ADHD load. Thus, during a developmental phase of “learning to speak”, where motor skills shape children’s learning environment and, in turn, behaviour and language learning^71^, children with a higher genetic predisposition for ADHD may be genetically inclined to express a larger, rather than smaller vocabulary size. In contrast, the polygenic relationship with ADHD reversed for receptive vocabulary in toddlerhood. Structural models showed that a genetic factor related to toddlerhood vocabulary measures, originating from expressive vocabulary size at 24 months of age, was inversely associated with ADHD symptom expression. These findings are consistent with known genetic associations of higher ADHD risk with lower child and adolescent verbal and cognitive abilities^22^. For phenotypes such as late-phase expressive vocabulary, sharing genetic influences with both early-phase expressive and late-phase receptive vocabulary, positive and negative genetic covariance patterns with ADHD may cancel, consistent with little evidence for a genome-wide genetic correlation of late-phase expressive vocabulary with ADHD.

Genetic and residual contributions to phenotypic correlations of vocabulary size are, however, complex. Positive genetic associations of early-phase expressive vocabulary with ADHD symptoms were found to be masked at the phenotypic level due to residual correlations with an opposite direction of effect. This rare violation of Cheverud’s conjecture^65^ suggests that residual sources of variation do not affect developmental pathways in the same way as genetic sources, and may implicate a protective effect of the caregiving environment, as observed for some behavioural traits in animals^72^. Thus, despite the validity of Cheverud’s conjecture in general^73^, additional research is required to characterise divergent genetic and residual association patterns in very young children.

This work has several strengths and limitations. Our work builds on a previous GWAS effort^9^ by increasing the number of studied children by ~50% and adopting a multivariate analysis approach to maximise statistical power while extending the studied phenotypic vocabulary spectrum. However, the power to detect single variant contributions of small effect (e.g. 0.1%) remained low (Supplemental Methods), especially for receptive vocabulary. The derived summary statistics did, however, capture a substantial fraction of phenotypic variance and had SNP-h^2^ Z-scores >4, suggesting validity for genetic covariance analyses^46^. Given limited data availability, this study focussed exclusively on European children and languages. Previous research has demonstrated the comparability of CDI measures across different European languages, but not yet non-European^74^. Finally, all measures were rank-transformed to harmonise vocabulary measures across different developmental stages and instruments. It is unlikely that this data transformation has affected the nature of our findings, as previous work demonstrated the robustness of identified phenotypic relationships using transformed vocabulary scores^8^. Future studies may increase the sample size further and boost study power through multivariate analysis of vocabulary with genetically related aspects of language, such as grammatical abilities^7,10^, preferably in genetically diverse populations.

In summary, there are at least two genetic factors contributing to vocabulary size during infancy and toddlerhood that match distinct polygenic association patterns with later-life traits. Our findings highlight the importance of studying genetic influences underlying early-life vocabulary acquisition to unravel aetiological processes that shape future behaviour and cognition.

## Acknowledgments

We are extremely grateful to all children, parents and caregivers for making this study possible. The project was embedded within the Early Genetics and Life Course Epidemiology (EAGLE) Consortium. Cohort-specific acknowledgments and funding information can be found in the Supplemental Methods. In addition, we would like to thank all cohorts and researchers that made their summary statistics available to us. This includes the Social Science Genetic Association Consortium (SSGAC), Early Growth Genetics (EGG) Consortium, Early Genetics and Life Course Epidemiology (EAGLE) Consortium, Psychiatric Genomics Consortium (PGC) and the Danish Lundbeck Foundation Initiative for Integrative Psychiatric Research (iPSYCH). EVe and BSTP had full access to all summary-statistic level data in the study and take responsibility for the integrity of the data and the accuracy of the data analysis. This article is available as preprint: https://www.biorxiv.org/content/10.1101/2022.06.01.494306.

Beate St Pourcain, Ellen Verhoef and the EAGLE working group indicate that the EAGLE working group is one of the named authors and that all members qualify for authorship. The EAGLE working group that contributed to this manuscript consists of the following members: Ole A. Andreassen^1,2,3^, Meike Bartels^4^, Dorret Boomsma^4^, Philip S. Dale^5^, Erik Ehli^6^, Dietmar Fernandez-Orth^7^, Mònica Guxens^7,8,9,10^, Christian Hakulinen^11^, Kathleen Mullan Harris^12,13^, Simon Haworth^14,15^, Vincent Jaddoe^16,17^, Liisa Keltikangas-Järvinen, Terho Lehtimäki^18,19,20^, Christel Middeldorp^21,22^, Josine L. Min^14,23^, Pashupati P. Mishra^18,19,20^, Pål Rasmus Njølstad^24,25^, Jordi Sunyer^7,8,9^, Ashley E. Tate^26^, Nicholas Timpson^14,23^, Camiel van der Laan^4^, Martine Vrijheid^7,8,9^, Eero Vuoksimaa^27^, Alyce Whipp^27^, Eivind Ystrom^28,29^, ACTION Consortium^30^, BIS investigator group^31^

1. NORMENT Centre, Institute of Clinical Medicine, University of Oslo, Oslo, Norway

2. Division of Mental Health and Addiction, Oslo University Hospital, Oslo, Norway

3. KG Jebsen Centre for Neurodevelopmental disorders, University of Oslo, Oslo, Norway

4. Netherlands Twin Register, dept Biological Psychology, Vrije Universiteit Amsterdam, the Netherlands

5. Speech & Hearing Sciences Department, University of New Mexico, Albuquerque, NM, USA

6. Avera Institute for Human Genetics, Sioux Falls, SD, USA

7. ISGlobal, Barcelona, Spain

8. Pompeu Fabra University, Barcelona, Spain

9. Spanish Consortium for Research on Epidemiology and Public Health (CIBERESP), Instituto de Salud Carlos III, Madrid, Spain

10. Department of Child and Adolescent Psychiatry/Psychology, Erasmus MC, University Medical Centre, Rotterdam, The Netherlands

11. Department of Psychology and Logopedics, University of Helsinki, Finland

12. Department of Sociology, University of North Carolina at Chapel Hill, NC, USA

13. Carolina Population Center, University of North Carolina at Chapel Hill, NC, USA

14. Medical Research Council Integrative Epidemiology Unit at the University of Bristol, Bristol, UK

15. Bristol Dental School, University of Bristol, Bristol, UK

16. Department of Pediatrics, Erasmus University Medical Center, Rotterdam, The Netherlands

17. Generation R Study Group, Erasmus University Medical Center Rotterdam

18. Department of Clinical Chemistry, Faculty of Medicine and Health Technology, Tampere University, Tampere, Finland

19. Finnish Cardiovascular Research Centre, Faculty of Medicine and Health Technology, Tampere University, Tampere, Finland

20. Department of Clinical Chemistry, Fimlab Laboratories, Tampere, Finland

21. University of Queensland, Child health Research Centre, Brisbane, Australia

22. Child and Youth Mental Health Service, Children’s Health Queensland Hospital and Health Service, Brisbane, Australia

23. Population Health Sciences, Bristol Medical School, University of Bristol, Bristol, UK

24. Department of Clinical Science, University of Bergen, NO-5020 Bergen, Norway

25. Children and Youth Clinic, Haukeland University Hospital, NO-5021 Bergen, Norway

26. Department of Medical Epidemiology and Biostatistics, Karolinska Institutet, Stockholm, Sweden

27. Institute for Molecular Medicine Finland (FIMM), HiLIFE, University of Helsinki, Finland

28. PROMENTA Research Center, Department of Psychology, University of Oslo, Oslo, Norway

29. Department of Mental Disorders, Norwegian Institute of Public Health, Oslo, Norway

30. Supplemental Methods, ACTION Consortium GWAMA authors and affiliations

31. Supplemental Methods, BIS investigator group authors and affiliations

EVe, EEi, FS, SEF and BSTP were funded by the Max Planck Society. TSA was supported by the Novo Nordisk Foundation Grant NNF18OC0052457. ATM is supported by the National Health and Medical Research Council. CS was supported by an Australian Government NHMRC Postgraduate Research Scholarship. EH receives funding from the National Health and Medical Research Council Australia, Australian Research Council, Medical Research Future Fund, and Tour de Cure. MG is funded by a Miguel Servet II fellowship (CPII18/00018) awarded by the Spanish Institute of Health Carlos III. We acknowledge support from the Spanish Ministry of Science and Innovation and the State Research Agency through the “Centro de Excelencia Severo Ochoa 2019-2023” Program (CEX2018-000806-S), and support from the Generalitat de Catalunya through the CERCA Program. SH receives support from the UK National Institute for Health Research through the academic clinical fellowship scheme. KR is supported by a Sir Henry Wellcome Postdoctoral Fellowship. CYS and JLM are supported by the UK Medical Research Council (MRC) Integrative Epidemiology Unit at the University of Bristol (MC_UU_00011/5). OAA is supported by KG Jebsen Stiftelsen, Research Council of Norway (#223273, 273291, 324252). EY receives support from the Research Council of Norway (grant numbers 262177; 288083). PRN was supported by grants from the European Research Council (AdG SELECTionPREDISPOSED #293574), the Bergen Research Foundation ("Utilizing the Mother and Child Cohort and the Medical Birth Registry for Better Health"), Stiftelsen Kristian Gerhard Jebsen (Translational Medical Center), the University of Bergen, the Research Council of Norway (FRIPRO grant #240413), the Western Norway Regional Health Authority (Strategic Fund “Personalized Medicine for Children and Adults”), the Novo Nordisk Foundation (grant #54741), and the Norwegian Diabetes Association. CAMC receives support from the European Union’s Horizon 2020 Research and Innovation Programme under grant agreement No 848158 (EarlyCause Project).

## References

1. Kennison SM. Introduction to Language Development. 1st ed. SAGA; 2014.

2. Clark EV. First Language Acquisition. Cambridge University Press; 2016.

3. Bergelson E, Swingley D. At 6-9 months, human infants know the meanings of many common nouns. Proc Natl Acad Sci USA. 2012;109(9):3253–3258. doi:10.1073/pnas.1113380109

4. Fenson L, Dale PS, Reznick JS, Bates E, Thal DJ, Pethick SJ. Variability in early communicative development. Monogr Soc Res Child Dev. 1994;59(5):1–173; discussion 174-85.

5. Goldfield BA, Reznick JS. Early lexical acquisition: rate, content, and the vocabulary spurt. J Child Lang. 1990;17(1):171–183.

6. Hoff E. Language Development. Cengage Learning; 2013.

7. Dionne G, Dale PS, Boivin M, Plomin R. Genetic Evidence for Bidirectional Effects of Early Lexical and Grammatical Development. Child Development. 2003;74(2):394–412. doi:10.1111/1467-8624.7402005

8. Verhoef E, Shapland CY, Fisher SE, Dale PS, St Pourcain B. The developmental genetic architecture of vocabulary skills during the first three years of life: Capturing emerging associations with later-life reading and cognition. PLOS Genetics. 2021;17(2):e1009144. doi:10.1371/journal.pgen.1009144

9. St Pourcain B, Cents RA, Whitehouse AJ, et al. Common variation near ROBO2 is associated with expressive vocabulary in infancy. Nature communications. 2014;5:4831. doi:10.1038/ncomms5831

10. Dale PS, Dionne G, Eley TC, Plomin R. Lexical and grammatical development: a behavioural genetic perspective. Journal of Child Language. 2000;27(03):619–642. doi:null

11. Hayiou-Thomas ME, Dale PS, Plomin R. The etiology of variation in language skills changes with development: a longitudinal twin study of language from 2 to 12 years. Dev Sci. 2012;15(2):233–249. doi:10.1111/j.1467-7687.2011.01119.x

12. Reznick JS, Corley R, Robinson J. A longitudinal twin study of intelligence in the second year. Monogr Soc Res Child Dev. 1997;62(1):i-vi, 1–154; discussion 155-60.

13. Harlaar N, Hayiou-Thomas ME, Dale PS, Plomin R. Why Do Preschool Language Abilities Correlate With Later Reading? A Twin Study. J Speech Lang Hear Res. 2008;51(3):688–705. doi:10.1044/1092-4388(2008/049)

14. Verhoef E, Shapland CY, Fisher SE, Dale PS, St Pourcain B. The developmental origins of genetic factors influencing language and literacy: Associations with early-childhood vocabulary. Journal of Child Psychology and Psychiatry. Published online September 14, 2020. doi:10.1111/jcpp.13327

15. Geurts HM, Embrechts M. Language profiles in ASD, SLI, and ADHD. J Autism Dev Disord. 2008;38(10):1931–1943. doi:10.1007/s10803-008-0587-1

16. Helland WA, Posserud MB, Helland T, Heimann M, Lundervold AJ. Language Impairments in Children With ADHD and in Children With Reading Disorder. Journal of Attention Disorders. Published online October 16, 2012:1087054712461530. doi:10.1177/1087054712461530

17. Germanò E, Gagliano A, Curatolo P. Comorbidity of ADHD and dyslexia. Dev Neuropsychol. 2010;35(5):475–493. doi:10.1080/87565641.2010.494748

18. Peyre H, Galera C, van der Waerden J, et al. Relationship between early language skills and the development of inattention/hyperactivity symptoms during the preschool period: Results of the EDEN mother-child cohort. BMC Psychiatry. 2016;16. doi:10.1186/s12888-016-1091-3

19. Martin NC, Levy F, Pieka J, Hay DA. A Genetic Study of Attention Deficit Hyperactivity Disorder, Conduct Disorder, Oppositional Defiant Disorder and Reading Disability: Aetiological overlaps and implications. International Journal of Disability, Development and Education. 2006;53(1):21–34. doi:10.1080/10349120500509992

20. Willcutt EG, Pennington BF, DeFries JC. Twin study of the etiology of comorbidity between reading disability and attention-deficit/hyperactivity disorder. Am J Med Genet. 2000;96(3):293–301. doi:10.1002/1096-8628(20000612)96:3<293::AID-AJMG12>3.0.CO;2-C

21. Willcutt EG, Pennington BF, Olson RK, DeFries JC. Understanding comorbidity: A twin study of reading disability and attention-deficit/hyperactivity disorder. Am J Med Genet. 2007;144B(6):709–714. doi:10.1002/ajmg.b.30310

22. Verhoef E, Demontis D, Burgess S, et al. Disentangling polygenic associations between attention-deficit/hyperactivity disorder, educational attainment, literacy and language. Translational Psychiatry. 2019;9(1):35. doi:10.1038/s41398-018-0324-2

23. Tager-Flusberg H, Paul R, Lord C, Volkmar FR, Klin A, Cohen D. Handbook of autism and pervasive developmental disorders. Published online 2005.

24. Ozonoff S, South M, Miller JN. DSM-IV-Defined Asperger Syndrome: Cognitive, Behavioral and Early History Differentiation from High-Functioning Autism. Autism. 2000;4(1):29–46. doi:10.1177/1362361300041003

25. Rocco I, Corso B, Bonati M, Minicuci N. Time of onset and/or diagnosis of ADHD in European children: a systematic review. BMC Psychiatry. 2021;21(1):575. doi:10.1186/s12888-021-03547-x

26. van ’t Hof M, Tisseur C, van Berckelear-Onnes I, et al. Age at autism spectrum disorder diagnosis: A systematic review and meta-analysis from 2012 to 2019. Autism. 2021;25(4):862–873. doi:10.1177/1362361320971107

27. Harris SR. Measuring head circumference: Update on infant microcephaly. Can Fam Physician. 2015;61(8):680–684.

28. Maunu J, Parkkola R, Rikalainen H, Lehtonen L, Haataja L, Lapinleimu H. Brain and Ventricles in Very Low Birth Weight Infants at Term: A Comparison Among Head Circumference, Ultrasound, and Magnetic Resonance Imaging. Pediatrics. 2009;123(2):617–626. doi:10.1542/peds.2007-3264

29. Bartholomeusz HH, Courchesne E, Karns CM. Relationship Between Head Circumference and Brain Volume in Healthy Normal Toddlers, Children, and Adults. Neuropediatrics. 2002;33(05):239–241. doi:10.1055/s-2002-36735

30. Middeldorp CM, Felix JF, Mahajan A, et al. The Early Growth Genetics (EGG) and EArly Genetics and Lifecourse Epidemiology (EAGLE) consortia: design, results and future prospects. Eur J Epidemiol. 2019;34(3):279–300. doi:10.1007/s10654-019-00502-9

31. Fenson L, Pethick S, Renda C, Cox JL, Dale PS, Reznick JS. Short-form versions of the MacArthur Communicative Development Inventories. Applied Psycholinguistics. 2000;21(01):95–116. doi:null

32. Zink I, Lejaegere M. N-CDI’s: korte vormen, Aanpassing en hernormering van de MacArthur Short Form Vocabulary Checklist van Fenson et al.

33. Reznick JS, Goldsmith L. A multiple form word production checklist for assessing early language. Journal of Child Language. 1989;16(01):91–100. doi:10.1017/S0305000900013453

34. Fenson L, Dale P, Reznick JS, et al. User’s Guide and Technical Manual for the MacArthur Communicative Development Inventories. Singular Publishing; 1993.

35. Bleses D, Vach W, Slott M, et al. The Danish Communicative Developmental Inventories: validity and main developmental trends. J Child Lang. 2008;35(3):651–669. doi:10.1017/S0305000907008574

36. Rescorla Leslie. The Language Development Survey. Journal of Speech and Hearing Disorders. 1989;54(4):587–599. doi:10.1044/jshd.5404.587

37. Marees AT, Kluiver H de, Stringer S, et al. A tutorial on conducting genome-wide association studies: Quality control and statistical analysis. International Journal of Methods in Psychiatric Research. 2018;27(2):e1608. doi:10.1002/mpr.1608

38. McCarthy S, Das S, Kretzschmar W, et al. A reference panel of 64,976 haplotypes for genotype imputation. Nat Genet. 2016;48(10):1279–1283. doi:10.1038/ng.3643

39. Winkler TW, Day FR, Croteau-Chonka DC, et al. Quality control and conduct of genome-wide association meta-analyses. Nat Protoc. 2014;9(5):1192–1212. doi:10.1038/nprot.2014.071

40. Willer CJ, Li Y, Abecasis GR. METAL: fast and efficient meta-analysis of genomewide association scans. Bioinformatics. 2010;26(17):2190–2191. doi:10.1093/bioinformatics/btq340

41. Turley P, Walters RK, Maghzian O, et al. Multi-trait analysis of genome-wide association summary statistics using MTAG. Nature Genetics. 2018;50(2):229–237. doi:10.1038/s41588-017-0009-4

42. Nyholt DR. A Simple Correction for Multiple Testing for Single-Nucleotide Polymorphisms in Linkage Disequilibrium with Each Other. Am J Hum Genet. 2004;74(4):765–769.

43. Watanabe K, Taskesen E, Bochoven A van, Posthuma D. Functional mapping and annotation of genetic associations with FUMA. Nature Communications. 2017;8(1):1826. doi:10.1038/s41467-017-01261-5

44. Ning Z, Pawitan Y, Shen X. High-definition likelihood inference of genetic correlations across human complex traits. Nat Genet. 2020;52(8):859–864. doi:10.1038/s41588-020-0653-y

45. Bulik-Sullivan BK, Loh PR, Finucane HK, et al. LD Score regression distinguishes confounding from polygenicity in genome-wide association studies. Nature genetics. 2015;47(3):291–295. doi:10.1038/ng.3211

46. Bulik-Sullivan B, Finucane HK, Anttila V, et al. An atlas of genetic correlations across human diseases and traits. Nat Genet. 2015;47(11):1236–1241. doi:10.1038/ng.3406

47. Savage JE, Jansen PR, Stringer S, et al. Genome-wide association meta-analysis in 269,867 individuals identifies new genetic and functional links to intelligence. Nature Genetics. 2018;50(7):912–919. doi:10.1038/s41588-018-0152-6

48. Lee JJ, Wedow R, Okbay A, et al. Gene discovery and polygenic prediction from a genome-wide association study of educational attainment in 1.1 million individuals. Nature Genetics. 2018;50(8):1112–1121. doi:10.1038/s41588-018-0147-3

49. Taal HR, St Pourcain B, Thiering E, et al. Common variants at 12q15 and 12q24 are associated with infant head circumference. Nat Genet. 2012;44(5):532–538. doi:10.1038/ng.2238

50. Haworth S, Shapland CY, Hayward C, et al. Low-frequency variation in TP53 has large effects on head circumference and intracranial volume. Nature Communications. 2019;10(1):357. doi:10.1038/s41467-018-07863-x

51. Ip HF, van der Laan CM, Krapohl EML, et al. Genetic association study of childhood aggression across raters, instruments, and age. Transl Psychiatry. 2021;11(1):1–9. doi:10.1038/s41398-021-01480-x

52. Jami ES, Hammerschlag AR, Ip HF, et al. Genome-wide association meta-analysis of childhood and adolescent internalising symptoms. Published online July 31, 2021:2020.09.11.20175026. Accessed August 12, 2021. https://www.medrxiv.org/content/10.1101/2020.09.11.20175026v2

53. Demontis D, Walters RK, Martin J, et al. Discovery of the first genome-wide significant risk loci for attention deficit/hyperactivity disorder. Nature Genetics. Published online November 26, 2018:1. doi:10.1038/s41588-018-0269-7

54. Grove J, Ripke S, Als TD, et al. Identification of common genetic risk variants for autism spectrum disorder. Nature Genetics. 2019;51(3):431. doi:10.1038/s41588-019-0344-8

55. Li J, Ji L. Adjusting multiple testing in multilocus analyses using the eigenvalues of a correlation matrix. Heredity (Edinb). 2005;95(3):221–227. doi:10.1038/sj.hdy.6800717

56. Grotzinger AD, Rhemtulla M, de Vlaming R, et al. Genomic structural equation modelling provides insights into the multivariate genetic architecture of complex traits. Nature Human Behaviour. 2019;3(5):513–525. doi:10.1038/s41562-019-0566-x

57. Boyd A, Golding J, Macleod J, et al. Cohort Profile: the ‘children of the 90s’--the index offspring of the Avon Longitudinal Study of Parents and Children. Int J Epidemiol. 2013;42(1):111–127. doi:10.1093/ije/dys064

58. Fraser A, Macdonald-Wallis C, Tilling K, et al. Cohort Profile: the Avon Longitudinal Study of Parents and Children: ALSPAC mothers cohort. Int J Epidemiol. 2013;42(1):97–110. doi:10.1093/ije/dys066

59. St Pourcain B, Eaves LJ, Ring SM, et al. Developmental changes within the genetic architecture of social communication behaviour: A multivariate study of genetic variance in unrelated individuals. Biological Psychiatry. 2017;83:598–606. doi:10.1016/j.biopsych.2017.09.020

60. Middeldorp CM, Hammerschlag AR, Ouwens KG, et al. A Genome-Wide Association Meta-Analysis of Attention-Deficit/Hyperactivity Disorder Symptoms in Population-Based Pediatric Cohorts. J Am Acad Child Adolesc Psychiatry. 2016;55(10):896–905.e6. doi:10.1016/j.jaac.2016.05.025

61. Neale M, Boker S, Xie G, Meas HHM. Mx: Statistical Modeling. 7th ed. Department of Psychiatry, Medical College of Virginia; 2006.

62. Falconer DS, Mackay TFC. Quantitative Genetics. Vol fourth. Pearson; 1996.

63. Leeuw CA de, Mooij JM, Heskes T, Posthuma D. MAGMA: Generalized Gene-Set Analysis of GWAS Data. PLOS Computational Biology. 2015;11(4):e1004219. doi:10.1371/journal.pcbi.1004219

64. Reilly S, Cook F, Bavin EL, et al. Cohort Profile: The Early Language in Victoria Study (ELVS). International Journal of Epidemiology. 2018;47(1):11–20. doi:10.1093/ije/dyx079

65. Cheverud JM. A Comparison of Genetic and Phenotypic Correlations. Evolution. 1988;42(5):958–968. doi:10.1111/j.1558-5646.1988.tb02514.x

66. Curtin S, Archer SL. Speech perception. In: Bavin EL, Naigles LR, eds. The Cambridge Handbook of Child Language. 2nd ed. Cambridge Handbooks in Language and Linguistics. Cambridge University Press; 2015:137–158. doi:10.1017/CBO9781316095829.007

67. Alcock K. The development of oral motor control and language. Down Syndrome Research and Practice. 2006;11(1):1–8. doi:10.3104/reports.310

68. Smith A, Goffman L, Stark RE. Speech Motor Development. Semin Speech Lang. 1995;16(2):87–99. doi:10.1055/s-2008-1064112

69. Hannigan LJ, Askeland RB, Ask H, et al. Developmental milestones in early childhood and genetic liability to neurodevelopmental disorders. Psychological Medicine. Published online September 21, 2021:1–9. doi:10.1017/S0033291721003330

70. Riglin L, Tobarra-Sanchez E, Stergiakouli E, et al. Early manifestations of genetic liability for ADHD, autism and schizophrenia at ages 18 and 24 months. JCPP Advances. 2022;2(3):e12093. doi:10.1002/jcv2.12093

71. Libertus K, Violi DA. Sit to Talk: Relation between Motor Skills and Language Development in Infancy. Front Psychol. 2016;7. doi:10.3389/fpsyg.2016.00475

72. Hadfield JD, Nutall A, Osorio D, Owens IPF. Testing the phenotypic gambit: phenotypic, genetic and environmental correlations of colour. Journal of Evolutionary Biology. 2007;20(2):549–557. doi:10.1111/j.1420-9101.2006.01262.x

73. Sodini SM, Kemper KE, Wray NR, Trzaskowski M. Comparison of Genotypic and Phenotypic Correlations: Cheverud’s Conjecture in Humans. Genetics. 2018;209(3):941–948. doi:10.1534/genetics.117.300630

74. Bleses D, Vach W, Slott M, et al. Early vocabulary development in Danish and other languages: a CDI-based comparison. J Child Lang. 2008;35(3):619–650. doi:10.1017/S0305000908008714

