## Supplemental Information for "Genome-wide analyses of vocabulary size in infancy and toddlerhood: associations with Attention-Deficit/Hyperactivity Disorder and cognition-related traits"

### Supplemental Methods

#### Study design

Expressive vocabulary scores were analysed as part of two developmental stages: an early phase (15-18 months, infancy) and a late phase (24-38 months, toddlerhood). The early phase reflects a developmental window during which children produce their first words, usually in isolation[1]. During the late phase, children start to use word combinations and more complex grammatical structures[2,3].

Receptive vocabulary scores were only studied for the late phase (24-38 months, toddlerhood), as parents tend to underestimate receptive vocabulary in children below the age of two years compared to direct assessment of child receptive language using a preferential looking task[4]. In addition, the availability of receptive vocabulary scores in infants is low and there was little evidence ($P$<0.05) for Single-Nucleotide Polymorphism (SNP) heritability at 15 months of age within the Avon Longitudinal Study of Parents and Children (ALSPAC)[5]. However, this does not exclude the existence of individual genetic variants that have an effect on vocabulary size.

#### Early vocabulary cohort descriptives

Avon Longitudinal Study Parents and Children: Pregnant women resident in Avon, UK with expected dates of delivery 1st April 1991 to 31st December 1992 were invited to take part in the Avon Longitudinal Study of Parents and Children (ALSPAC)[6,7]. The initial number of pregnancies enrolled is 14,541 (for these at least one questionnaire has been returned or a "Children in Focus" clinic had been attended by 19/07/99). Of these initial pregnancies, there was a total of 14,676 foetuses, resulting in 14,062 live births and 13,988 children who were alive at 1 year of age.

When the oldest children were approximately 7 years of age, an attempt was made to bolster the initial sample with eligible cases who had failed to join the study originally. As a result, when considering variables collected from the age of seven onwards (and potentially abstracted from obstetric notes) there are data available for more than the 14,541 pregnancies mentioned above. The number of new pregnancies not in the initial sample (known as Phase I enrolment) that are currently represented on the built files and reflecting enrolment status at the age of 24 is 913 (456, 262 and 195 recruited during Phases II, III and IV respectively), resulting in an additional 913 children being enrolled. The phases of enrolment are described in more detail in the cohort profile paper and its update. The total sample size for analyses using any data collected after the age of seven is therefore 15,454 pregnancies, resulting in 15,589 foetuses. Of these 14,901 were alive at 1 year of age.

A 10% sample of the ALSPAC cohort, known as the Children in Focus (CiF) group, attended clinics at the University of Bristol at various time intervals between 4 to 61 months of age. The CiF group were chosen at random from the last 6 months of ALSPAC births (1,432 families attended at least one clinic). Excluded were those mothers who had moved out of the area or were lost to follow-up, and those partaking in another study of infant development in Avon.

Ethical approval for the study was obtained from the ALSPAC Ethics and Law Committee and the Local Research Ethics Committees. Consent for biological samples has been collected in accordance with the Human Tissue Act (2004). Informed consent for the use of data collected via questionnaires and clinics was obtained from participants following the recommendations of the ALSPAC Ethics and Law Committee at the time.

Please note that the study website contains details of all the data that are available through a fully searchable data dictionary and variable search tool (http://www.bristol.ac.uk/alspac/researchers/our-data/).

Barwon Infant Study: The Barwon Infant Study (BIS) is a prospective birth cohort with antenatal recruitment based in the Barwon region of Victoria, southwest of Melbourne, Australia. From June 2010 to June 2013, a birth cohort of 1,074 mother–infant pairs (10 sets of twins) was recruited using an unselected antenatal sampling frame[8]. Eligibility criteria, population characteristics, and measurement details have been described previously[8]. In brief, women were recruited prior to 28 weeks' gestation between 2010 and 2013, and infant exclusion criteria were: (1) delivery before 32 weeks, (2) serious neonatal illness, (3) major congenital malformation or genetic disease and (4) family having moved out of the Barwon Statistical Division by the time of birth. Informed consent was obtained from pregnant mothers. Mother–infant pairs were reviewed at regular intervals at birth; 4 weeks; 3, 6, 9, 12 and 18 months; 2 years and 4 years; with review at 7-10 years in progress. Further details of the cohort are available through the Barwon Infant Study cohort website (https://www.barwoninfantstudy.org.au/index.cfm). A short-form adaption of the MacArthur Communicative Development Inventories: Words and Sentences was completed for a subsample of N=431 children at the 2-year-old review alongside other neurodevelopmental assessments. The current analysis was restricted to those children determined to be of European ancestry based on their genetic data (N=383). Ethics approval for the study was obtained from the Barwon Health Human Research Ethics Committee (HREC 10/24).

Copenhagen Prospective Studies on Asthma in Childhood: The Copenhagen Prospective Studies on Asthma in Childhood is a clinical study with multiple cohorts (COPSAC2000 and COPSAC2010). The COPSAC2010 cohort is a population-based prospective mother-child cohort comprising 700 children born to unselected mothers (during 2009-10) from Zealand, Denmark. The cohort was enrolled at age 1 week and attended the research clinic for clinical examinations at ages 1, 3, 6, 9, 12, 18, 24, 30 and 36 month and yearly hereafter till age 8 years.

Language development was assessed with the Danish version of The McArthur Bates Communicative Development Inventory (CDI) through questionnaires filled by parents as described elsewhere[9]. Genotyping and imputation for the COPSAC2010 cohort has been described elsewhere[10]. The Ethics Committee for Copenhagen and the Danish Data Protection Agency approved this study.

Early Language in Victoria Study: The Early Language in Victoria Study (ELVS) is a longitudinal, community cohort study that comprehensively tracked the language development of a large group of children (>1,900) born in the surrounds of Melbourne, Victoria (Australia). ELVS was designed to fill knowledge gaps about language development and factors that predict later outcomes, including service costs and health-related quality of life. The children were recruited between 8 and 10 months of age and have been followed across key developmental transitions, including infancy and early childhood, middle childhood to adolescence and most recently into early adulthood. Data were obtained via multi-source informants, direct assessment and linkage to nationally acquired academic achievement data. ELVS operates from Melbourne, Australia, and is funded by the Australian National Health and Medical Research Council (#237106, #436958, #1041947). Ethical approval has been obtained from the Royal Children's Hospital Human Research Ethics Committee (27078/33195). A number of ELVS sub-studies investigate different areas of communication development, including stuttering, autism and bilingual language development. Full details of the study and the ages at which data were collected are available on the Lifecourse website (https://lifecourse.melbournechildrens.com/cohorts/elvs/) and in Reilly et al (2018)[11].

Generation R Study: The Generation R Study (GenR) is a prospective cohort study from fetal life onwards that included pregnant women living in Rotterdam, the Netherlands, with an expected delivery date between April 2002 and January 2006 (N=9,778). The main aim of this study is to identify early environmental and genetic factors that affect growth, health and development[12]. The Generation R Study is multidisciplinary, and both prenatal and postnatal measures have included multiple domains of growth, health and development. Rotterdam is an ethnically diverse city and this is reflected in the Generation R participants. Of the enrolled mothers, 42% were of non-Dutch ethnic background, largely made by mothers from Surinamese (9%), Turkish (7%) and Moroccan (3%) background[12,13]. Data have been collected in children up until the mean age of 13 years, with current on-going data collection at mean age 17 years.

Study protocols were approved by the local ethics committee, and written informed consent and assent was obtained from all parents and children.

The Growing Up in Australia: Longitudinal Study of Australian Children study: The Growing Up in Australia: Longitudinal Study of Australian Children study (LSAC) includes prospective birth (B) and kinder (K, not further considered in the EAGLE consortium) cohorts that aimed to be broadly representative of the Australian population[14]. The B-cohort recruited 5,107 0-1-year-olds in 2004, with continued in-home follow-up every 2 years. The Child Health CheckPoint module was one-off physical health and biomarkers assessment, nested between LSAC's 10-11 year and 12-13 year waves, for 1,874 B-cohort families, at either an assessment centre or home visit, and including biospecimen collection for DNA extraction[15]. The Australian Institute of Family Studies (AIFS) Ethics Committee approved each wave of LSAC. The AIFS Ethics Committee (14-26) and Melbourne's Royal Children's Hospital Human Research Ethics Committee (33225D) approved the CheckPoint wave. Written informed consent and assent was obtained from all parents for LSAC, and from all parents and children for the CheckPoint.

The Raine Study: The Raine Study is a prospective pregnancy cohort where 2,900 mothers (Gen1) were recruited between 1989 and 1991[16,17]. Recruitment took place at Western Australia's major perinatal centre, King Edward Memorial Hospital, and nearby private practices. Women who had sufficient English language skills, an expectation to deliver at King Edward Memorial Hospital, and an intention to reside in Western Australia to allow for future follow-up of their child (Gen2) were eligible for the study.

The Raine Study is known as one of the largest successful prospective cohorts richly phenotyped at multiple time points over pregnancy, infancy, childhood, adolescence, and young adulthood. The mothers (Gen1) completed questionnaires regarding their children (Gen2), and the children (Gen2) had physical examinations at ages 1, 2, 3, 6, 8, 10, 14, 17, 20 and 22 years. Ethics approval for the original pregnancy cohort and subsequent follow-ups were granted by the Human Research Ethics Committee of King Edward

Memorial Hospital, Princess Margaret Hospital, the University of Western Australia, and the Health Department of Western Australia.

Twins Early Development Study: The Twins Early Development Study (TEDS) is a longitudinal twin study that recruited over 16,000 twin pairs born between 1994 and 1996 in England and Wales through national birth records[18]. More than 10,000 of these families are still involved in the study. TEDS was, and still is, a representative sample of the population in England and Wales. Rich cognitive and behavioural data have been collected from the twins from infancy to emerging adulthood, with data collection at ages 2, 3, 4, 7, 8, 9, 10, 12, 14, 16, 18, 19 and 21, enabling longitudinal genetically sensitive study designs. Data have been collected from the twins themselves (including extensive web-based cognitive testing), and from their parents and teachers. Genotyped DNA data are available for 10,346 individuals (who are unrelated except for 3,320 dizygotic co-twins). Ethical approval was received from King's College London Research Ethics Committee (Reference number PNM/09/10-104).

### Early vocabulary assessment

The CDIs were developed to assess language and communication development in young children[19], whereas the Language Development Survey (LDS) aims to identify children with language delays[20]. Vocabulary scores were primarily assessed in English (ALSPAC, BIS, LSAC, the Raine Study and TEDS), but also in Danish (COPSAC, Danish adaptation of the MacArthur CDI:Words & Sentences[21]) and Dutch (GenR, N-CDI-2A[22]). Research has shown that children follow similar patterns of language acquisition across different languages[23] and that CDI vocabulary assessments are comparable across different cultures, including English, Dutch and Danish[24].

Early-phase expressive vocabulary: During the early phase (15-18 months, infancy), expressive vocabulary size was assessed using an abbreviated form of the MacArthur CDI:Words & Gestures[25] in the

ALSPAC cohort (15 months: N=6,741). Early-phase expressive vocabulary was defined by this instrument as the total number of words a child could "say and understand" and thus jointly represents expressive and receptive vocabulary. Within GenR (18 months: N=2,058), expressive vocabulary was assessed using a Dutch adaptation of the short-form version of the MacArthur CDI (N-CDI-2A)[22]. This form included the response "say" in addition to "say and understand", so early-phase expressive vocabulary was defined as the number of words that fell into either of these categories.

Late-phase expressive vocabulary: During the late phase (24-38 months, toddlerhood), expressive vocabulary size was assessed with an abbreviated version of the MacArthur CDI:Words & Sentences[19] in ALSPAC (24 months: N=6,208; 38 months: N=6,291), the corresponding Danish adaptation[21] in COPSAC (24 months: N=487), and using the LDS[20] in GenR (31 months: N=1,825) and the Raine Study (26 months: N=980). Adapted forms of the MacArthur CDI[26,27] (MCDI) were used to assess expressive vocabulary in BIS (30 months: N=383), LSAC (34 months: N=1,134), and TEDS (24 months: N=5,515). For CDI vocabulary assessments[19,21,26,27], late-phase expressive vocabulary was defined as the number of words that fell within the categories "says" and "says and understands". For LDS vocabulary assessments[20], late-phase expressive vocabulary was defined as the total number of words spontaneously produced by a child from a given list of words. The LDS and CDI have high concurrent validity, with a correlation of 0.95 on total vocabulary scores[28].

Late-phase receptive vocabulary: Late-phase receptive vocabulary scores were only available in ALSPAC (38 months: N=6,291) and assessed using an abbreviated form of the MacArthur CDI:Words & Sentences[19] (Table S1). Late-phase receptive vocabulary score was defined as the number of words a child could understand, regardless of whether they also produced the word, and encoded as "understand" plus "say and understand".

Single-variant association analysis

Genome-wide association study per cohort: Within each cohort, vocabulary scores were adjusted for age, sex, $age^2$ and their interaction effects, as well as ancestry-informative principal components (that differed by cohort) and other study-specific covariates, such as genotyping array and/or batch, defined by the local genome-wide association study (GWAS) analyst. Vocabulary scores were rank-transformed to achieve normality and allow for comparisons of genetic effects across different psychological instruments.

Single-Nucleotide Polymorphism (SNP)-vocabulary associations were estimated within each cohort using a linear regression of rank-transformed residuals on posterior genotype probability using SNPTEST[29], Proabel[30] and GEMMA[31] software, assuming an additive genetic model, except for the LSAC cohort. For LSAC, a linear regression of rank-transformed residualised vocabulary scores on bestguess genotypes was performed with PLINK 1.9[32], using imputed markers (INFO>0.3) as posterior genotype probability data were not available (Table S2). GWAS analyses of twin samples were performed using GEMMA[31] following a linear mixed-model approach. This method accounts for relatedness among individuals using a genetic-relationship matrix (GRM) derived from high-quality, directly genotyped markers. GRM off-diagonal elements ≥0.05 captured relatedness for closely related individuals[33,34], while other elements of the GRM were set to zero.

Quality control at summary statistic level: GWAS summary statistics from all cohorts underwent extensive quality control using the EasyQC R package[35] (v9.2): variants that had low (i) imputation quality (INFO<0.6 for SNPTEST, PLINK and GEMMA association analyses and INFO<0.5 for Proabel association analyses), (ii) minor allele count (MAC≤10), or (iii) effect allele frequency (EAF≤0.005 or EAF≥0.995) were excluded. In addition, marker names were harmonised, and alleles were aligned against Haplotype Reference Consortium (HRC) r1.1 reference data. Variants with missing or mismatching alleles were dropped, as well as all insertions/deletions, duplicate SNPs and multi-allelic SNPs. Finally, variants with an EAF that deviated >0.2 from the frequency in the HRC r1.1. reference data were excluded. All association analyses were applied with genomic control[36] for variant discovery and without genomic control for follow-up analyses.

Single-trait meta-GWAS (stage I): As part of analysis stage I (Figure 1), fixed-effect meta-analyses were carried out for early-phase expressive vocabulary using METAL software[37]. This approach includes a meta-analysis across effect size estimates reported by each individual cohort, weighted by the inverse of the corresponding standard error[37]. As late-phase expressive vocabulary included longitudinal assessments of the same ALSPAC children at 24 and 38 months (Table S1), a fixed-effect meta-analysis was carried out excluding ALSPAC expressive vocabulary at 38 months to ensure the independence of GWAS summary statistics across cohorts. The derived METAL output was then jointly analysed with the GWAS results for ALSPAC expressive vocabulary at 38 months using multi-trait analysis of genome-wide association (MTAG)[38]. This method exploits genetic relationships among traits and provides a generalised inverse-variance-weighted meta-analysis estimate by integrating GWAS summary statistics across different traits while allowing for overlapping samples[38]. As late-phase receptive vocabulary scores were only available for ALSPAC, no meta-analysis was performed.

Multi-trait meta-GWAS (stage II): As part of analysis stage II, multi-trait meta-analyses were performed with MTAG[38] combining vocabulary summary statistics with moderate-to-strong genetic correlations ($r_g \geq 0.65$) to increase statistical power (Figure 1). Late-phase expressive vocabulary, the most powerful measure, was included as the outcome in all multi-trait meta-analyses.

Sensitivity analyses for MTAG: To assure robustness of our findings, sensitivity analyses were conducted for all meta-analyses performed with MTAG. MTAG analyses combining low-powered traits (mean $\chi^2$ statistic <1.02), such as ALSPAC expressive vocabulary at 38 months and late-phase receptive vocabulary, may lead to bias and an increased false discovery rate[38]. Therefore, MTAG-derived estimates for each meta-analysis estimates were compared against fixed-effect meta-analysis estimates for late-phase expressive vocabulary (the most powerful fixed-effect meta-analysis, see above). For each meta-analysis, we compared beta coefficients and standard errors across all SNPs ($N_{SNPs}$=7,343,861-7,355,069), as well as across a subset of highly-associated SNPs ($P<5x10^{-6}$, $N_{SNPs}$=37).

Power analysis: The single-variant association analyses with the largest sample size (late-phase expressive and receptive vocabulary, stage II) had 99% power to detect association with a genetic variant explaining 0.3% of the trait variance (assuming an additive model and an increaser allele frequency of 0.1, with complete LD with marker and genetic risk variant)[39]. However, power to detect variants with smaller contributions to the trait variance was still only modest (e.g. 27% power to detect a genetic variant explaining 0.1% of the trait variance)[39].

### FUMA analyses

Gene-based GWASs: Gene-based GWASs were conducted with Multi-marker Analysis of GenoMic Annotation (MAGMA, v1.08) according to a SNP-wide mean model[40], as implemented within FUMA software[41] (v1.3.6a). SNPs were mapped to genes using positional mapping based on the 1000 Genomes Phase 3 European reference panel (release 20130502) and a 0kb window, consistent with default MAGMA settings. SNPs were mapped to ≤18,896 protein-coding genes. Assuming 2.38 independent vocabulary measures, estimated with a Matrix Spectral Decomposition (matSpD)[42] of bivariate genetic correlations (see Main), this resulted in a genome-wide gene-based significance threshold of $1.11\times10^{-6}$ (0.05/18,896/2.38). Gene-based GWAS results subsequently served as input for gene-set and gene-property analyses (see below).

Gene-set analyses: MAGMA-based gene-set analyses[40] (v1.08) were performed as implemented within FUMA software[41] (v1.3.6a). This competitive test was conditioned on gene size, gene density, and the inverse of the mean minor allele count in the gene[40]. Association was investigated with up to 4,527 gene ontology (GO) biological pathways that were derived from MsigDB[43] v7.0 and contained between 10 and 200 genes to avoid bias related to gene-set size[44]. The multiple-testing-adjusted threshold was defined at $P<4.64\times10^{-6}$ (0.05/4,527/2.38).

Gene-property analyses: MAGMA[40] gene-property analyses were performed in FUMA[41] (v1.3.6a) to assess whether common genetic variation related to vocabulary was enriched for expression in certain tissues and/or developmental periods of interest. For these analyses, gene expression data from 30 broad tissue types and 54 specific tissues derived from the GTEx v8 RNA-sequencing database[45] were obtained, as well as gene expression data for 29 different age groupings and 11 developmental stages from BrainSpan[46]. The multiple-testing-adjusted threshold was defined at $P<1.69\text{x}10^{-4}$, accounting for the total number of gene expression data sets and independent vocabulary measures investigated (0.05/124/2.38).

### High-Definition Likelihood SNP-heritability and genetic correlation analyses

SNP-heritability (SNP-$h^2$) and bivariate genetic correlations ($r_g$), as captured by GWAS summary statistics, were estimated using High-Definition Likelihood[47] (HDL). HDL is a full likelihood-based method that naturally extends the Linkage Disequilibrium Score (LDSC) regression formula by including non-diagonal elements of Z-score covariance matrices. Compared to LDSC, HDL estimates SNP-$h^2$ and $r_g$ with increased accuracy[47]. HDL-SNP-$h^2$ analyses were conducted using the pre-computed reference panel for European-ancestry populations based on 1,029,876 high-quality UK Biobank imputed HapMap3 SNPs if >99% of them were available, following HDL recommendations[47]. Otherwise, a reference panel based on 769,306 high-quality UK Biobank imputed HapMap2 SNPs was used. These reference panels were created previously, and details, including quality control, are described elsewhere[47]. Eigenvalues and eigenvectors were derived by HDL, selecting values that resulted in the most stable heritability estimate.

We investigated evidence for genetic correlation based on GWAS summary statistics for vocabulary size, created as part of stage I, and several heritable health-, cognition-, and behaviour-related outcomes (SNP-$h^2$ $P$<0.05 and SNP-$h^2$ Z-score >4, Table S7). All traits included in HDL-$r_g$ analyses had sufficient SNP overlap (>99%) with either the HapMap2 or HapMap3 reference panel. Genome-wide summary statistics for the studied health-, cognition-, and behaviour-related traits are described below in brief, while more detailed information can be found in the original studies:

<u>Intelligence:</u> GWAS summary statistics on intelligence across the lifespan (5-98 years, N=279,930) were obtained from the Complex Traits Genetics lab[48] (https://ctg.cncr.nl/documents/p1651/SavageJansen_IntMeta_sumstats.zip). Each cohort assessed intelligence with different instruments that were re-defined to index a common latent factor of general intelligence[48].

<u>Educational attainment:</u> GWAS summary statistics on years-of-schooling (>30 years, N=766,345 excluding 23andMe) were obtained from the Social Science Genetic Association Consortium[49] (https://www.dropbox.com/s/ho58e9jmytmpaf8/GWAS_EA_excl23andMe.txt?dl=0). Educational attainment (EA) was coded according to the International Standard Classification of Education (1997) scale[49] and analysed as a quantitative variable defined as an individual's years of schooling[49].

<u>Infant head circumference:</u> GWAS summary statistics on infant head circumference (6-30 months, N=10,768) were obtained from the Early Growth Genetics Consortium[50] (http://egg-consortium.org/HC/EGG_HC_DISCOVERY.v2.txt.gz). Head circumference was measured from the occipital protuberance to the forehead, using a flexible, non-stretching measure tape following standardized procedures[50].

<u>Childhood head circumference:</u> GWAS summary statistics on childhood head circumference[51] (6-9 years, N=10,600) were retrieved via Dr. Beate St Pourcain. Head circumference

was measured with a measuring tape at the widest horizontal circumference in the majority of participants[51].

Childhood aggressive behaviour: GWAS summary statistics on childhood aggressive behaviour[52] (1.5-18 years, N=151,741) were obtained via Prof. dr. Dorret Boomsma. Aggression was assessed on continuous scales, with higher scores indicating more aggressive behaviour, using mothers, fathers, teachers, and self-report based on 26 different instruments[52].

Childhood internalising symptoms: GWAS summary statistics on childhood internalising symptoms[53] (3-18 years, N=64,641) were retrieved via Prof. dr. C.M. Middeldorp. In the absence of diagnostic data, internalising symptoms were dimensionally measured and positively scored on continuous scales, with higher scores indicating more internalising symptoms, based on different raters and instruments[53].

Attention-Deficit/Hyperactivity Disorder: GWAS summary statistics on Attention-Deficit/Hyperactivity Disorder (ADHD) were accessed through the Danish Lundbeck Foundation Initiative for Integrative Psychiatric Research (iPSYCH) and Psychiatric Genetics Consortium (PGC)[54]. Analyses were restricted to individuals of European ancestry (https://ipsych.dk/en/research/downloads/data-download-agreement-adhd-european-ancestry-gwas-june-2017/). ADHD cases in iPSYCH were identified from a national research register and diagnosed by psychiatrists at a psychiatric hospital according to ICD10[55] (F90.0) and identified using the Danish Psychiatric Central Research Register[56]. ADHD cases in PGC were primarily diagnosed with the Diagnostic and Statistical Manual of Mental Disorders (DSM-III[57], DSM-IV[57], DSM-IV-TR[57]) or the International Classification of Diseases (ICD-10[55]).

Autism Spectrum Disorder: GWAS summary statistics on Autism Spectrum Disorder (ASD) were accessed through iPSYCH and PGC[58] (https://ipsych.dk/en/research/downloads/data-download-agreement-ipsych-pgc-asd-nov2017/). ASD cases were diagnosed according to ICD-10[55] and identified using the Danish Psychiatric Central Research Register[56]. Registry-based ASD diagnoses were validated previously[54,58]. Controls were randomly selected from the same nationwide birth cohort and did not have a diagnosis of ASD or ADHD, or moderate-severe mental retardation (F71-F79)[54,58,59]. In addition, data from five family-based trio samples of European ancestry from the PGC were included[60], which based an ASD diagnosis on the Autism Diagnostic Interview-Revised[61], the Autism Diagnostic Observation Schedule[62], and/or the Autism Screening Questionnaire[63]. The sample only included individuals of European ancestry.

Power to detect a genetic correlation between single-trait vocabulary data (stage I) and educational attainment was calculated via an online tool (https://eagenetics.shinyapps.io/power_website/) following Dudbridge et al[64]. Sample sizes of 8,800, 19,300, 6,300 and 766,300 were used for early-phase expressive vocabulary, late-phase expressive vocabulary, late-phase receptive vocabulary and educational attainment, respectively. SNP-$h^2$ estimates for these traits are reported in Table S5 and S7. As all traits are continuous, sample and population prevalence were set to one. Alpha was set to the multiple-testing-adjusted significance threshold of $5.32 \times 10^{-3}$.

### Polygenic scoring analyses

Due to limited data availability, only late-phase expressive vocabulary could be studied with out-of-sample prediction. Here, we studied phenotypic and genome-wide genetic data from ELVS[65] as the target sample and single-trait late-phase expressive vocabulary summary statistics (stage I) as the discovery sample.

Phenotypic data: In ELVS, late-phase expressive vocabulary was assessed using an adapted version of the MacArthur CDI:Words & Sentences at 24 months of age (N=639, Table S1). ELVS CDI vocabulary scores were adjusted for age, sex, age$^2$, their interaction effects, and the first two principal components, and rank-transformed.

Genetic data: ELVS individuals were genotyped using the Infinium Global Screening Array, and standard quality control procedures were applied[66]. To assure high-quality genetic data, variants with a call rate <0.98, Hardy-Weinberg Equilibrium $<1x10^{-6}$ or minor allele frequency <0.01 were excluded. Individuals were excluded based on a call rate <0.98, a non-European genetic ancestry, or relatedness with other participants (IBD >0.125). Cleaned genotype data were imputed using the Michigan Imputation Server (https://imputationserver.sph.umich.edu/index.html) against the HRC r1.1 reference panel[67]. For polygenic scoring analyses, allele counts for SNPs with an INFO score >0.8, genotyping probability >0.9 in 95% of the individuals and minor allele frequency >0.005 were transformed into bestguess genotypes ($N_{SNPs}$=6,675,600).

Polygenic scoring: Posterior SNP effect size estimates from late-phase expressive vocabulary summary statistics (stage I) were estimated using PRS-CS[68]: a Bayesian-based approach that adjusts SNP effect sizes for LD by applying a continuous-shrinkage parameter. Following the default settings: the global shrinkage parameter was learned from the data using a fully Bayesian approach, parameters a and b in the gamma-gamma prior were set to 1 and 0.5, respectively, and 1,000 Markov Chain Monte Carlo iterations were performed, with 500 burn-in iterations and a Markov chain thinning factor of 5. The 1000 Genomes phase 3 European reference panel provided by the authors was used as LD reference panel. Individual-level polygenic scores in ELVS were created with PLINK[32] (v1.9b3w) and Z-standardised. Finally, vocabulary measures were regressed on polygenic scores in ELVS using ordinary least square (OLS) regression (R:stats library, Rv4.1.0), and the phenotypic variance explained was assessed with the regression $R^2$.

Power calculation: Power to detect a genetic relationship was derived following Dudbridge et al.[64] via an online tool: https://eagenetics.shinyapps.io/power_website/. Parameters included for the discovery trait (meta-GWAS stage I, late-phase expressive vocabulary) were a sample size of 19,300, SNP-$h^2$ of 0.08, and a population and sample prevalence of 1. For the target trait (ELVS, late-phase expressive vocabulary) these parameters were set at N=600, SNP-$h^2$=0.08, and a population and sample prevalence of 1. The alpha was set at 0.05.

### Structural equation modelling

To obtain insight into the developmentally changing genetic correlation pattern of ADHD symptoms with vocabulary size across infancy and toddlerhood, we applied genetic-relationship-matrix structural equation modelling[69] (GRM-SEM) using the grmsem R package (v1.1.2) and individual-level data from the Avon Longitudinal Study of Parents And Children (ALSPAC) cohort. Note that it was not possible to fit an informative summary statistic-based SEM model with HDL-based Genomic SEM[70], as this requires >99% overlap with the same reference panel (HapMap2 or HapMap3) across all summary statistics of interest and sufficient SNP-$h^2$. In our case, not all traits of interest had sufficient overlap with HapMap3 SNPs and using the Hapmap2 reference panel led to a drop in SNP-$h^2$.

Vocabulary data: Data on expressive vocabulary size at 15 months (early-phase expressive vocabulary), 24 months and 38 months (late-phase expressive vocabulary), as well as receptive vocabulary size at 38 months (late-phase receptive vocabulary) from ALSPAC children were analysed in a similar way to the presented meta-GWASs (Table S1, Table S2), except for stricter filtering on relatedness (IBD <0.05) due to methodological assumptions[69]. This resulted in individual-level genotype and phenotype data for 6,524, 6,014, 6,092, and 6,092 children for expressive vocabulary at 15 months, expressive vocabulary at 24 months, expressive vocabulary at 38 months and receptive vocabulary at 38 months, respectively.

ADHD symptom score data: ADHD symptom scores in ALSPAC (showing strong genetic correlations with ADHD status in case-control analyses[71]) were assessed with the hyperactivity scale of the Strengths and Difficulties Questionnaire[72] (SDQ) at 7, 10, 12, 13 and 17 years of age using mother reports and at 8 and 11 years using teacher reports (Table S3). Only pro-rated scores were selected for the current study.

ADHD symptom scores were adjusted for age, sex, their interaction effects and the first two principal components and then rank-transformed.

SNP-$h^2$ analyses: Across ADHD SDQ symptom scores, SNP-$h^2$ was estimated using Genome-based restricted maximum likelihood (GREML) analyses[73,74], as implemented in Genome-wide Complex Trait Analysis (GCTA) software[75], based on a GRM including directly genotyped SNPs only. The two ADHD symptom scores with the highest SNP-$h^2$ Z-score based on mother- and teacher-report (Table S3) were selected for subsequent analyses (see below), as disorder-related genetic effects captured by parent and teacher reports may differ[76].

GRM-SEM analyses: A Cholesky decomposition was fitted to the data using genetic-relationship-matrix structural equation modelling[69] (GRM-SEM) with the grmsem R package (v1.1.2). A Cholesky decomposition is a saturated model that decomposes the phenotypic variance into as many latent genetic (A) and residuals (E) factors as there are observed variables, without any restrictions on the structure[77]. Subsequently, genetic ($r_g$) and residual ($r_e$) covariance and correlations were estimated as outlined by theory[78]. The Cholesky model included all four early vocabulary measures available in ALSPAC (expressive vocabulary at 15 months, 24 months and 38 months, as well as receptive vocabulary at 38 months) in addition to ADHD symptom scores at 8 and 13 years (in this order). Cholesky decompositions were fitted, allowing for missing data.

### Cohort-specific acknowledgements

ALSPAC: We are extremely grateful to all the families who took part in this study, the midwives for their help in recruiting them, and the whole ALSPAC team, which includes interviewers, computer and laboratory technicians, clerical workers, research scientists, volunteers, managers, receptionists and nurses. The UK Medical Research Council and Wellcome (Grant ref: 217065/Z/19/Z) and the University of Bristol provide core support for ALSPAC. This publication is the work of the authors and they will serve as guarantors for the contents of this paper. A comprehensive list of grants funding is available on the ALSPAC website (http://www.bristol.ac.uk/alspac/external/documents/grant-acknowledgements.pdf). GWAS data were generated by Sample Logistics and Genotyping Facilities at Wellcome Sanger Institute and LabCorp (Laboratory Corporation of America) using support from 23andMe.

BIS: We thank the BIS participants for the generous contribution they have made to this project. We also thank current and past staff for their efforts in recruiting and maintaining the cohort and in obtaining and processing the data and biospecimens. The establishment work and infrastructure for the BIS was provided by the Murdoch Children's Research Institute, Deakin University and Barwon Health. Subsequent funding was secured from the National Health and Medical Research Council of Australia (NHMRC), the Australian Research Council, the Jack Brockhoff Foundation, the Scobie & Claire Mackinnon Trust, the Shane O'Brien Memorial Asthma Foundation, the Our Women's Our Children's Fund Raising Committee Barwon Health, The Shepherd Foundation, the Rotary Club of Geelong, the Ilhan Food Allergy Foundation, GMHBA Limited, Vanguard Investments Australia Ltd, the Percy Baxter Charitable Trust, Perpetual Trustees, the Gandel Foundation, the Gwenyth Raymond Trust, and the Minderoo Foundation. In kind support was provided by the Cotton On Foundation and CreativeForce. Research at the Murdoch Children's Research Institute is supported by the Victorian Government's Operational Infrastructure Support Programme.

The BIS Investigator Group also consists of: Peter Vuillermin, Anne-Louise Ponsonby, Peter Sly, David Burgner, Fiona Collier, John Carlin, Lawrence Gray, Len Harrison, Martin O'Hely, Sarath Ranganathan, and Amy Loughman. We thank Terry Dwyer and Katie Allen for their past work as foundation investigators and John Carlin for statistical advice in the Barwon Infant Study.

COPSAC: We express our deepest gratitude to the children and families of the COPSAC cohort study for all their support and commitment. We acknowledge and appreciate the unique efforts of the COPSAC research team. All funding received by COPSAC is listed on www.copsac.com. The Lundbeck Foundation (Grant no R16-A1694); The Ministry of Health (Grant no 903516); Danish Council for Strategic Research (Grant no 0603-00280B) and The Capital Region Research Foundation have provided core support to the COPSAC research center.

ELVS: We thank those who have dedicated their time and expertise to ELVS over the past 13 years, including Ms. Eileen Cini, Dr. Angela Pezic, Dr. Laura Conway, Prof. Cristina McKean, Prof. James Law, Prof. John Carlin, Prof. Fiona Mensah, Prof. Melissa Wake, Prof. Edith Bavin, Assoc. Prof Obioha Ukoumunne, Prof. Margot Prior and the families who have generously given their time to participating in ELVS. The research was supported by the National Health and Medical Research Council (grant numbers 237106, 436958, 1041947, 1023493).

GenR: The Generation R Study is conducted by the Erasmus Medical Center in close collaboration with the Erasmus University Rotterdam, Faculty of Social Sciences, the Municipal Health Service Rotterdam area, and the Stichting Trombosedienst and Artsenlaboratorium Rijnmond (STAR), Rotterdam. We gratefully acknowledge the contribution of general practitioners, hospitals, midwives and pharmacies in Rotterdam. The Generation R Study is made possible by financial support from: Erasmus Medical Center, Rotterdam, and the Netherlands Organization for Health Research and Development (ZonMw).

LSAC: The study is conducted in partnership between the Department of Social Services, the Australian Institute of Family Studies, and the Australian Bureau of Statistics. Its biophysical module, the Child Health CheckPoint, was led by researchers at the Murdoch Children's Research Institute and collaborators. The findings and views reported are those of the authors. We thank the LSAC and CheckPoint (Child Health CheckPoint) study participants, staff, and students for their contributions. The CheckPoint module was supported by the National Health and Medical Research Council (NHMRC) of Australia [1041352, 1109355]; the Royal Children's Hospital Foundation [2014-241]; the Murdoch Children's Research Institute (MCRI); The University of Melbourne, the National Heart Foundation of Australia

[100660]; Financial Markets Foundation for Children [2014-055, 2016-310]; the Victoria Deaf Education Institute; and MBIE Catalyst grant (The New Zealand-Australia Life Course Collaboration on Genes, Environment, Nutrition and Obesity (GENO); UOAX1611).

The Raine Study: This study was supported by the National Health and Medical Research Council of Australia [grant numbers 572613, 403981 and 1059711] and the Canadian Institutes of Health Research [grant number MOP-82893]. The authors are grateful to the Raine Study participants and their families, and to the Raine Study research staff for cohort coordination and data collection. The authors gratefully acknowledge the NH&MRC for their long term funding to the study over the last 30 years and also the following institutes for providing funding for Core Management of the Raine Study: The University of Western Australia (UWA) , Curtin University, the Raine Medical Research Foundation, the UWA Faculty of Medicine, Dentistry and Health Sciences, the Telethon Kids Institute, the Women's and Infant's Research Foundation (King Edward Memorial Hospital), Murdoch University, The University of Notre Dame (Australia), and Edith Cowan University. The authors gratefully acknowledge the assistance of the Western Australian DNA Bank (National Health and Medical Research Council of Australia National Enabling Facility). Andrew Whitehouse is funded by Senior Research Fellowship from the NHMRC (1077966). We would also like to acknowledge the Raine Study participants for their ongoing participation in the study, and the Raine Study Team for study co-ordination and data collection. This work was supported by resources provided by the Pawsey Supercomputing Centre with funding from the Australian Government and Government of Western Australia.

TEDS: We gratefully acknowledge the ongoing contribution of the participants in TEDS and their families. TEDS is supported by a programme grant to RP from the UK Medical Research Council Program Grant MR/V012878/1 (and previously Grant MR/M021475/1), with additional support from the US National Institutes of Health (Grant No. AG046938). The research leading to these results has also received funding from the European Research Council under the European Union's Seventh Framework Programme

(FP7/2007-2013)/ grant agreement no. 602768. This research was funded in whole, or in part by the Wellcome Trust (213514/Z/18/Z). For the purpose of open access, the author has applied a CC BY public copyright licence to any Author Accepted Manuscript version arising from this submission.

### ACTION Consortium GWAMA authors and affiliations

Ip, Hill F.[1,] van der Laan, Camiel M.[1,2], Krapohl, Eva M. L.[3], Brikell, Isabell[4], Sánchez-Mora, Cristina[5,6,7], Nolte, Ilja M.[8], St Pourcain, Beate[9,10,11], Bolhuis, Koen[12], Palviainen, Teemu[13], Zafarmand, Hadi[14,15], Colodro-Conde, Lucía[15], Gordon, Scott[16], Zayats, Tetyana[17,18,19], Aliev, Fazil[20,21], Jiang, Chang[22,23], Wang, Carol A.[24], Saunders, Gretchen[25], Karhunen, Ville[26], Hammerschlag, Anke R.[1,27,28], Adkins, Daniel E.[29,30], Border, Richard[31,32,33], Peterson, Roseann E.[34], Prinz, Joseph A.[35], Thiering, Elisabeth[36,37], Seppälä, Ilkka[38], Vilor-Tejedor, Natàlia[39-43], Ahluwalia, Tarunveer S.[44,45], Day, Felix R.[46], Hottenga, Jouke-Jan[1], Allegrini, Andrea G.[3], Rimfeld, Kaili[3], Chen, Qi[4], Lu, Yi[4], Martin, Joanna[4,47], Soler Artigas, María[5,6,7], Rovira, Paula[5,6,7], Bosch, Rosa[5,6,48], Español, Gemma[5], Ramos Quiroga, Josep Antoni[5,6,7,48], Neumann, Alexander[12,49], Ensink, Judith[50,51], Grasby, Katrina[16], Morosoli, José J.[15], Tong, Xiaoran[22,23], Marrington, Shelby[52], Middeldorp, Christel[27,1,53], Scott, James G.[52,54,55], Vinkhuyzen, Anna[56], Shabalin, Andrey A.[30], Corley, Robin[31,57], Evans, Luke M.[31,57], Sugden, Karen[58,35], Alemany, Silvia[39,40,41], Sass, Lærke[44], Vinding, Rebecca[44], Ruth, Kate[59], Tyrrell, Jess[59], Ehli, Erik A.[60], Hagenbeek, Fiona A.[1], De Zeeuw, Eveline[1], Van Beijsterveldt, Toos C.E.M.[1], Larsson, Henrik[4,61], Snieder, Harold[8], Verhulst, Frank C.[12,62], Amin, Najaf[63], Whipp, Alyce M.[13], Korhonen, Tellervo[13], Vuoksimaa, Eero[13], Rose, Richard J.[64], Uitterlinden, André G.[63,65,66], Heath, Andrew C.[67], Madden, Pamela[67], Haavik, Jan[17,68], Harris, Jennifer R.[69], Helgeland, Øyvind[70], Johansson, Stefan[17,71], Knudsen, Gun Peggy S.[69], Njolstad, Pal Rasmus[72], Lu, Qing[22,23], Rodriguez, Alina[26,73], Henders, Anjali K.[56], Mamun, Abdullah[74], Najman, Jackob M.[52], Brown, Sandy[75], Hopfer, Christian[76], Krauter, Kenneth[77], Reynolds, Chandra[78], Smolen, Andrew[31], Stallings, Michael[31,32], Wadsworth, Sally[31], Wall, Tamara L.[75], Silberg, Judy L.[79,34], Miller, Allison[80], Keltikangas-Järvinen, Liisa[81], Hakulinen, Christian[81], Pulkki-Råback, Laura[81], Havdahl, Alexandra[82,83], Magnus, Per[84],

Raitakari, Olli T.[85,86,87], Perry, John R.B.[46], Llop, Sabrina[88,89], Lopez-Espinosa, Maria-Jose[88,89,90], Bønnelykke, Klaus[44], Bisgaard, Hans[44], Sunyer, Jordi[39,40,41,91], Lehtimäki, Terho[38], Arseneault, Louise[92], Standl, Marie[36], Heinrich, Joachim[36,93,94], Boden, Joseph[95], Pearson, John[96], Horwood, L John[95], Kennedy, Martin[97], Poulton, Richie[98], Eaves, Lindon J.[79,34], Maes, Hermine H.[79,34,99], Hewitt, John[31,32], Copeland, William E.[100], Costello, Elizabeth J.[101], Williams, Gail M.[52], Wray, Naomi[56,102], Järvelin, Marjo-Riitta[26,103], McGue, Matt[25], Iacono, William[25], Caspi, Avshalom[58,92,104,35], Moffitt, Terrie E.[58,92,104,35], Whitehouse, Andrew[105], Pennell, Craig E.[24], Klump, Kelly L.[106], Burt, S. Alexandra[106], Dick, Danielle M.[20,107,108], Reichborn-Kjennerud, Ted[83,109], Martin, Nicholas G.[15], Medland, Sarah E.[15], Vrijkotte, Tanja[14], Kaprio, Jaakko[110,13], Tiemeier, Henning[12,111], Davey Smith, George[9,112], Hartman, Catharina A.[113], Oldehinkel, Albertine J.[113], Casas, Miquel[5,6,7,48], Ribasés, Marta[5,6,7], Lichtenstein, Paul[4], Lundström, Sebastian[114,115], Plomin, Robert[3], Bartels, Meike[1,28], Nivard, Michel G.[1], Boomsma, Dorret .I.[1,27]

1. Department of Biological Psychology, Vrije Universiteit Amsterdam, Amsterdam, The Netherlands
2. The Netherlands Institute for the Study of Crime and Law Enforcement
3. Institute of Psychiatry, Psychology and Neuroscience, King's College London, United Kingdom
4. Department of Medical Epidemiology and Biostatistics, Karolinska Institutet, Stockholm, Sweden
5. Department of Psychiatry, Hospital Universitari Vall d'Hebron, Barcelona, Spain
6. Biomedical Network Research Centre on Mental Health (CIBERSAM), Instituto de Salud Carlos III, Barcelona, Spain
7. Psychiatric Genetics Unit, Group of Psychiatry, Mental Health and Addiction, Vall d'Hebron Research Institute (VHIR), Universitat Autònoma de Barcelona, Barcelona, Spain
8. Department of Epidemiology, University of Groningen, University Medical Center Groningen, Groningen, The Netherlands
9. MRC Integrative Epidemiology Unit, University of Bristol, Bristol, UK

10. Max Planck Institute for Psycholinguistics, The Netherlands
11. Donders Institute for Brain, Cognition and Behaviour, Radboud University, The Netherlands
12. Department of Child and Adolescent Psychiatry/Psychology, Erasmus University Medical Center, Rotterdam, Netherlands
13. Institute for Molecular Medicine FIMM, HiLife, University of Helsinki, Helsinki, Finland
14. Department of Public Health, Amsterdam Public Health Research Institute, Amsterdam UMC, location Academic Medical Center, University of Amsterdam, Amsterdam, The Netherlands
15. Department of Clinical Epidemiology, Biostatistics and Bioinformatics, Amsterdam Public Health Research Institute, Amsterdam UMC, location Academic Medical Center, University of Amsterdam, Amsterdam, The Netherlands
16. QIMR Berghofer Medical Research Institute, Brisbane, Australia
17. Department of Biomedicine, University of Bergen, Norway
18. Analytic and Translational Genetics Unit, Department of Medicine, Massachusetts General Hospital and Harvard Medical School, Boston, Massachusetts, USA
19. Stanley Center for Psychiatric Research, Broad Institute of MIT and Harvard, Cambridge, Massachusetts, USA
20. Department of Psychology, Virginia Commonwealth University, USA
21. Faculty of Business, Karabuk University, Turkey
22. Department of Epidemiology and Biostatistics, Michigan State University, East Lansing, USA
23. Department of Biostatistics, University of Florida, Gainesville, USA
24. School of Medicine and Public Health, Faculty of Medicine and Health, The University of Newcastle
25. Department of Psychology, University of Minnesota, USA
26. Department of Epidemiology and Biostatistics, MRC-PHE Centre for Environment and Health, School of Public Health, Imperial College London, London, W2 1PG, United Kingdom

27. Child Health Research Centre, the University of Queensland, Brisbane, QLD, Australia
28. Amsterdam Public Health Research Institute, Amsterdam, The Netherlands
29. Department of Sociology, College of Social and Behavioral Science, University of Utah
30. Department of Psychiatry, School of Medicine, University of Utah
31. Institute for Behavioral Genetics, University of Colorado Boulder, Colorado, USA
32. Department of Psychology and Neuroscience, University of Colorado Boulder, Colorado, USA
33. Department of Applied Mathematics, University of Colorado Boulder, Colorado, USA
34. Department of Psychiatry, Virginia Institute for Psychiatric and Behavioral Genetics, Virginia Commonwealth University
35. Center for Genomic and Computational Biology, Duke University, Durham, NC, USA
36. Institute of Epidemiology, Helmholtz Zentrum München - German Research Center for Environmental Health, Neuherberg, Germany
37. Ludwig-Maximilians-University of Munich, Dr. von Hauner Children's Hospital, Division of Metabolic Diseases and Nutritional Medicine, Munich, Germany
38. Department of Clinical Chemistry, Fimlab Laboratories, and Finnish Cardiovascular Research Center - Tampere, Faculty of Medicine and Health Technology, Tampere University, Tampere 33520, Finland
39. ISGlobal, Barcelona Institute for Global Health, Barcelona, Spain
40. Universitat Pompeu Fabra (UPF), Barcelona, Spain
41. CIBER Epidemiología y Salud Pública (CIBERESP), Spain
42. Centre for Genomic Regulation (CRG),The Barcelona Institute of Science and Technology, Barcelona, Spain
43. BarcelonaBeta Brain Research Center, Pasqual Maragall Foundation (FPM), Barcelona, Spain

44. COPSAC, Copenhagen Prospective Studies on Asthma in Childhood, Herlev and Gentofte Hospital, University of Copenhagen, Copenhagen, Denmark
45. Steno Diabetes Center Copenhagen, Gentofte, Denmark
46. MRC Epidemiology Unit, University of Cambridge School of Clinical Medicine, Institute of Metabolic Science, Cambridge Biomedical Campus, Cambridge, UK
47. MRC Centre for Neuropsychiatric Genetics and Genomics, Division of Psychological Medicine and Clinical Neurosciences, Cardiff University, Cardiff, UK
48. Department of Psychiatry and Legal Medicine, Universitat Autònoma de Barcelona, Barcelona, Spain
49. Lady Davis Institute for Medical Research, Jewish General Hospital, Montreal, Qc, Canada
50. Department of Child and Adolescent Psychiatry, Academic Medical Center, Amsterdam, The Netherlands
51. De Bascule, Academic centre for Child and Adolescent Psychiatry, Amsterdam, The Netherlands
52. School of Public Health, Faculty of Medicine, The University of Queensland, Herston 4006, Australia
53. Children's Health Queensland Hospital and Health Service, Child and Youth Mental Health Service, Brisbane, QLD, Australia
54. Metro North Mental Health, University of Queensland, QLD, Australia
55. Queensland Centre for Mental Health Research, QLD, Australia
56. Institute for Molecular Bioscience, University of Queensland, QLD, Australia
57. Department of Ecology and Evolutionary Biology, University of Colorado Boulder, Colorado, USA
58. Department of Psychology and Neuroscience and Center for Genomic and Computational Biology, Duke University, Durham, NC, USA
59. Genetics of Complex Traits, University of Exeter Medical School, Royal Devon & Exeter Hospital, Exeter, EX2 5DW, UK

60. Avera Institute for Human Genetics, Sioux Falls, South Dakota, USA
61. School of Medical Sciences, Orebro University, Orebro, Sweden
62. Child and Adolescent Mental Health Centre, Mental Health Services Capital Region, Research Unit, Copenhagen University Hospital, Copenhagen, Denmark
63. Department of Epidemiology, Erasmus University Medical Center, Rotterdam, The Netherlands
64. Department of Psychological and Brain Sciences, Indiana University, Bloomington, IN, USA
65. Department of Internal Medicine, Erasmus University Medical Center, Rotterdam, The Netherlands
66. Netherlands Genomics Initiative (NGI)-sponsored Netherlands Consortium for Healthy Aging (NCHA), Leiden, The Netherlands
67. Washington University, St Louis MO, USA
68. Division of Psychiatry, Haukeland University Hospital, Norway
69. Division of Health Data and Digitalisation, The Norwegian Institute of Public Health, Oslo, Norway
70. Department of Genetics and Bioinformatics, Division of Health Data and Digitalization, The Norwegian Institute of Public Health
71. K.G. Jebsen Centre for Neuropsychiatric Disorders, Department of Clinical Science, University of Bergen, Norway
72. Department of Clinical Science, University of Bergen, Norway
73. School of Psychology, University of Lincoln, Lincolnshire, LN5 7AY, United Kingdom
74. Institute for Social Science Research, University of Queensland, Long Pocket 4068, Australia
75. Department of Psychiatry, University of California, San Diego
76. University of Colorado School of Medicine, Aurora, Colorado, USA
77. Department of Molecular, Cellular, and Developmental Biology, University of Colorado Boulder, USA
78. Department of Psychology, University of California Riverside, Riverside, CA, USA

79. Department of Human & Molecular Genetics, Virginia Institute for Psychiatric and Behavioral Genetics, Virginia Commonwealth University
80. Department of Pathology and Biomedical Science, and Carney Centre for Pharmacogenomics, University of Otago Christchurch, New Zealand
81. Department of Psychology and Logopedics, Faculty of Medicine, University of Helsinki, Finland
82. Nic Waals Institute, Lovisenberg Diaconal Hospital, Oslo, Norway
83. Department of Mental Disorders, Norwegian Institute of Public Health, Oslo, Norway
84. Centre for Fertility and Health, Norwegian Institute of Public Health, Oslo, Norway
85. Centre for Population Health Research, University of Turku and Turku University Hospital
86. Research Centre of Applied and Preventive Cardiovascular Medicine, University of Turku
87. Department of Clinical Physiology and Nuclear Medicine, Turku University Hospital, Turku, Finland
88. Epidemiology and Environmental Health Joint Research Unit, FISABIO-Universitat Jaume I-Universitat de València, Valencia, Spain
89. Spanish Consortium for Research on Epidemiology and Public Health (CIBERESP), Madrid, Spain
90. Faculty of Nursing and Chiropody, Universitat de València, Valencia, Spain
91. IMIM (Hospital del Mar Medical Research Institute), Barcelona, Spain
92. Department of Psychiatry and Behavioral Sciences, Duke University School of Medicine, Durham, NC, USA
93. Institute and Outpatient Clinic for Occupational, Social and Environmental Medicine, University of Munich Medical Center, Ludwig-Maximilians-Universität München, Munich, Germany
94. Allergy and Lung Health Unit, Melbourne School of Population and Global Health, University of Melbourne, Melbourne, Australia
95. Christchurch Health and Development Study, Department of Psychological Medicine, University of Otago Christchurch, New Zealand

96. Biostatistics and Computational Biology Unit, Department of Pathology and Biomedical Science, University of Otago Christchurch, New Zealand

97. Department of Pathology and Biomedical Science, and Carney Centre for Pharmacogenomics, University of Otago Christchurch, New Zealand

98. Dunedin Multidisciplinary Health and Development Research Unit, University of Otago, Dunedin, New Zealand

99. Massey Cancer Center, Virginia Commonwealth University

100. Department of Psychiatry, College of Medicine, University of Vermont

101. Department of Psychiatry, School of Medicine, Duke University

102. Queensland Brain Institute, Institute for Molecular Bioscience, University of Queensland, St Lucia 4072, Australia

103. Center for Life Course Health Research, Faculty of Medicine, University of Oulu, PO Box 8000, FI-90014 Oulun yliopisto, Finland

104. Social, Genetic, and Developmental Psychiatry Research Centre, Institute of Psychiatry, Psychology, and Neuroscience, King's College London, UK

105. Telethon Kids Institute, The University of Western Australia, Perth, Western Australia, Australia

106. Department of Psychology, Michigan State University, East Lansing, USA

107. Department of Human and Molecular Genetics, Virginia Commonwealth University, Richmond, VA, USA

108. College Behavioral and Emotional Health Institute, Virginia Commonwealth University, Richmond, VA, USA

109. Institute of Clinical Medicine, University of Oslo, Oslo, Norway

110. Department of Public Health, Medical Faculty, University of Helsinki, Helsinki, Finland

111. Department of Social and Behavioral Science, Harvard TH Chand School of Public Health, Boston, USA

112. Population Health Sciences, Bristol Medical School, University of Bristol, Bristol, UK

113. Department of Psychiatry, University of Groningen, University Medical Center Groningen, Groningen, The Netherlands

114. Gillberg Neuropsychiatry Centre, University of Gothenburg, Gothenburg, Sweden

115. Sweden Centre for Ethics, Law and Mental Health, University of Gothenburg, Gothenburg, Sweden

### BIS investigator group authors and affiliations

Vuillermin, P.[1], Ponsonby, A-L.[2,3,4], Tang, M.[3,4,5], Burgner, D.[4,5,6], Harrison, L.[7], Gray, L.[8,9], Sly, P.[8,10]

1. Deakin University, Waurne Ponds, Geelong, VIC, Australia
2. The Florey Institute of Neuroscience and Mental Health
3. Royal Children's Hospital, Melbourne
4. Murdoch Children's Research Institute, University of Melbourne, Parkville, VIC, Australia
5. Department of Paediatrics, University of Melbourne
6. Department of Paediatrics, Monash University
7. Walter and Eliza Hall Institute of Medical Research
8. Faculty of Health, Deakin University, Geelong, VIC Australia
9. Child Health Research Unit at Barwon Health
10. Children's Health and Environment Program, Child Health Research Centre, The University of Queensland

### Supplemental tables

#### Table S1: Overview of participating cohorts

| Analysis | Cohort | Measure | Psychological Instrument | Raw vocabulary score (SD) | Age (SD) in months | N individuals (males) |
|---|---|---|---|---|---|---|
| meta-GWAS | ALSPAC | Early-phase EV | MacArthur CDI:Words & Gestures[a] | 14.34(17.84) | 15.42(0.98) | 6,741(3,445) |
| | | Late-phase EV | MacArthur CDI:Words & Sentences[a] | 64.10(35.20) | 24.39(1.02) | 6,208(3,197) |
| | | | | 113.28(17.5) | 38.48(1.19) | 6,291(3,226) |
| | | Late-phase RV | MacArthur CDI:Words & Sentences[a] | 109.66(23.78) | 38.48(1.19) | 6,291(3,226) |
| | BIS | Late-phase EV | MCDI:UKSF | 78.31(20.08) | 29.62(1.92) | 383(210) |
| | COPSAC | Late-phase EV | MacArthur CDI:Words & Sentences | 253.00(158.12) | 24.18(0.28) | 487(256) |
| | GenR | Early-phase EV | N-CDI-2A | 17.51(17.05) | 18.36(0.96) | 2,058(1,054) |
| | | Late-phase EV | LDS | 245.86(53.67) | 31.32(2.04) | 1,825(937) |
| | LSAC | Late-phase EV | MCDI | 56.95(23.60) | 33.51(2.51) | 1,134(558) |
| | Raine Study | Late-phase EV | LDS | 185.60(83.44) | 25.52(1.74) | 980(504) |
| | TEDS | Late-phase EV | MCDI | 48.66(24.79) | 24.48(1.20) | 5,515(2,665) |
| PGS | ELVS | Late-phase EV | MacArthur CDI:Words & Sentences | 269.82(157.16) | 24.13(0.29) | 639(314) |

a. abbreviated form

Expressive and receptive vocabulary size were assessed between 15 and 38 months of age using parental questionnaires. Data from seven independent cohorts were utilised in meta-genome-wide association analyses. Polygenic scoring analyses were performed using an eighth independent sample. For each cohort, the psychological instrument, mean raw vocabulary score and age, with corresponding standard errors, as well as sample size are reported. The instruments for early vocabulary assessment are described in detail in Supplemental Methods.

Abbreviations: ALSPAC, Avon Longitudinal Study of Parents and Children; BIS, Barwon Infant Study; CDI, Communicative Development Inventory; COPSAC, Copenhagen Prospective Studies on Asthma in Childhood; ELVS; Early Language in Victoria Study; EV, expressive vocabulary; GenR, Generation Rotterdam; LDS; Language Development Survey; LSAC, Longitudinal Study of Australian Children; MCDI, MacArthur Communicative Development Inventory; PGS, polygenic scoring; RV, receptive vocabulary; TEDS, Twins Early Development Study; UKSF, UK short form

Table S2: Overview of genotyping, imputation and analysis software

| | Cohort | ALSPAC | BIS | COPSAC | GenR | LSAC | Raine Study | TEDs |
|---|---|---|---|---|---|---|---|---|
| Genotyping platform | | Illumina HumanHap550 quad chip | Illumina Global Screening Array platform | Illumina Infinium HumanOmni ExomeExpress | Illumina 610K | Illumina Infinium® Global Screening Array-24 v1.0 | Illuminia Human660W Quad BeadChip | AffymetrixGeneChip (Affy) 6.0 and HumanOmniExpres sExome-8v1.2 (OEE) |
| Quality control | MAF | ≥0.01 | ≥0.01 | ≥0.01 | ≥0.01 | ≥0.01 | ≥0.01 | ≥0.01 |
| | SNP call rate | ≥0.99 | ≥0.99 | ≥0.95 | ≥0.95 | ≥0.95 | ≥0.95 | ≥0.99 |
| | HWE | $\geq 5x10^{-7}$ | $\geq 5x10^{-7}$ | $\geq 1x10^{-5}$ | $\geq 1x10^{-5}$ | $\geq 5x10^{-7}$ | $\geq 1x10^{-6}$ | $\geq 1x10^{-4}$ |
| | Individual call rate | ≥0.97 | ≥0.97 | ≥0.95 | ≥0.95 | ≥0.97 | ≥0.97 | ≥0.99 |
| | N SNPs genotyped | 440,476 | 451,479 | 566,755 | 477,033 | 468,271 | 517,183 | Affy: 608,517<br>OEE: 502,434 |
| Imputation | Platform | Sanger Imputation Server | Sanger Imputation Server | Sanger Imputation Server | Michigan Imputation Server | Sanger Imputation Server | Michigan Imputation Server | Sanger Imputation Server |
| | Reference panel | HRC (r1.1) | HRC (r1.1) | HRC (r1.1) | HRC (r1.1) | HRC (r1.1) | HRC (r1.1) | HRC (r1.1) |
| GWAS | Analysis software | SNPTEST | SNPTEST | SNPTEST | SNPTEST | PLINK | Proabel | GEMMA |
| | N SNPs after QC | Early-phase EV: 8,663,580<br><br>Late-phase EV: 8,665,928 (24m) 8,667,217 (38m)<br><br>Late-phase RV: 8,667,217 | 7,244,741 | 7,795,895 | Early-phase EV: 8,610,574<br><br>Late-phase EV: 8,607,086 | 7,518,913 | 8,654,834 | 8,293,360 |

Genotyping data for each cohort were obtained using high-density SNP arrays. Standard genomic quality control procedures were applied and genotypes were imputed against the HRC r1.1. reference panel[67] using either the Sanger imputation server (EAGLE2[79] v2.0.5 and PBWT[80] software, https://imputation.sanger.ac.uk/) or Michigan imputation server[81] (Minimac 3/4 and Shapeit v2.r790, https://imputationserver.sph.umich.edu/). Association analyses within cohorts of unrelated individuals were performed using SNPTEST[29], PLINK[32] and Proabel[30]. Genome-wide association analyses of related individuals were performed using GEMMA[31].

Abbreviations: ALSPAC, Avon Longitudinal Study of Parents and Children; BIS, Barwon Infant Study; COPSAC, Copenhagen Prospective Studies on Asthma in Childhood; EV, expressive vocabulary; GenR, Generation Rotterdam; GWAS, genome-wide association study; HRC, Haplotype Reference Consortium; HWE, Hardy-Weinberg Equilibrium; LSAC, Longitudinal Study of Australian Children; m, months; MAF, minor allele frequency; RV, receptive vocabulary; SNPs, Single-Nucleotide Polymorphism; TEDS, Twins Early Development Study

Table S3: ADHD symptom scores in the Avon Longitudinal Study of Parents and Children

| Trait | Reporter | Raw trait score (SD) | Age (SD) in years | N individuals (males) | SNP-h$^2$ (SE) | SNP-h$^2$ Z-score |
|---|---|---|---|---|---|---|
| ADHD symptoms | Mother | 3.33(2.35) | 6.79(0.11) | 5,348(2,782) | 0.08(0.06) | 1.26 |
| | | 2.90(2.23) | 9.65(0.12) | 5,516(2,797) | 0.08(0.06) | 1.19 |
| | | 2.74(2.21) | 11.72(0.13) | 5,110(2,546) | 0.19(0.07) | 2.77 |
| | | 2.90(2.22) | 13.16(0.18) | 4,929(2,458) | 0.22(0.07) | 3.16 |
| | | 2.53(2.11) | 16.84(0.36) | 4,061(1,979) | 0.09(0.09) | 1.08 |
| | Teacher | 2.48(2.62) | 8.33(0.31) | 3,572(1,801) | 0.27(0.10) | 2.86 |
| | | 2.15(2.60) | 11.16(0.33) | 4,254(2,132) | 0.16(0.08) | 2.05 |

ADHD symptom scores for unrelated children (IBD<0.05) were obtained from the Avon Longitudinal Study of Parents and Children. ADHD symptoms were assessed with the hyperactivity scale of the Strengths and Difficulties Questionnaire by mothers or teachers at various ages. For each assessment, mean raw trait score and age, with corresponding standard errors, as well as sample size are reported. SNP-h$^2$ estimates were derived using Genome-based restricted maximum likelihood (GREML) analyses[73,74], as implemented in Genome-wide Complex Trait Analysis (GCTA) software[75], based on a genetic-relationship matrix including directly genotyped SNPs only. SNP-h$^2$ Z-scores were calculated by dividing SNP-h$^2$ by its standard error.

Abbreviations: ADHD, Attention-Deficit/Hyperactivity Disorder; SNP, Single-Nucleotide Polymorphism

Table S4: MAGMA gene-set and gene-property analyses

| Analysis | | Early-phase EV | Late phase EV | Late phase RV |
|---|---|---|---|---|
| Gene-set | 4,527 GO biological pathways | $P \geq 5x10^{-5}$ | $P \geq 2x10^{-5}$ | $P \geq 8x10^{-6}$ |
| Gene-property | GTEx v8 30 broad tissue types | $P \geq 0.09$ | $P \geq 0.05$ | $P \geq 0.04$ |
| | GTEx v8 54 specific tissue types | $P \geq 0.14$ | $P \geq 0.13$ | $P \geq 0.02$ |
| | BrainSpan 29 ages | $P \geq 0.06$ | $P \geq 4x10^{-4}$ | $P \geq 0.12$ |
| | BrainSpan 11 developmental periods | $P \geq 0.07$ | $P \geq 0.11$ | $P \geq 0.12$ |

MAGMA[40] gene-set and gene-property analyses were performed in FUMA[41] (v1.3.6a). Association with 4,527 GO biological pathways containing between 10 and 200 genes was tested and the significance threshold adjusted for multiple-testing was determined at $P \leq 4.64x10^{-6}$, correcting for both the number of gene-sets tested and the estimated number of independent traits studied. Gene-property analyses were based on gene expression data from 30 broad tissue types and 54 specific tissue types from the GTEx v8 RNA sequencing database[45]. In addition, gene expression data from 29 different age groupings and 11 developmental stages from the BrainSpan database[46] were utilised. The lowest *P*-value obtained for each association analyses is reported. Gene-property analyses were considered significant if they passed a multiple-testing-adjusted *P*-value threshold of $1.45x10^{-4}$.

Abbreviations: EV, expressive vocabulary; GO, gene ontology; RV, receptive vocabulary

Table S5: SNP-heritability of vocabulary size based on summary statistics

| | | | HDL | | | LDSC |
|---|---|---|---|---|---|---|
| **meta-GWAS** | **Trait** | **$N_{ind}$** | **SNP-$h^2$(SE)** | **Z-score** | ***P*** | **SNP-$h^2$(SE)** |
| Stage I | Early-phase EV | 8,799 | 0.24(0.02) | 8.64 | $<1x10^{-10}$ | 0.12(0.05) |
| | Late phase EV | 19,296ǂ | 0.08(0.01) | 5.53 | $3x10^{-8}$ | 0.09(0.03) |
| | Late phase RV | 6,291 | 0.20(0.04) | 5.21 | $2x10^{-7}$ | 0.09(0.07) |
| Stage II | EV | 22,104ǂ | 0.10(0.01) | 6.91 | $<1x10^{-10}$ | 0.10(0.03) |
| | Late-phase ERV | 23,466ǂ | 0.07(0.01) | 5.00 | $5x10^{-7}$ | 0.11(0.03) |

SNP-heritability (SNP-$h^2$) was estimated for both single- (stage I) and multi-trait (stage II) vocabulary summary statistics using High-Definition Likelihood[47] (HDL), and LD Score Regression[82] (LDSC) for comparison. SNP-heritability, corresponding standard error and *P*-value were estimated with HDL using a HapMap3 reference panel. SNP-$h^2$ Z-scores were calculated by dividing SNP-$h^2$ by its standard error. For comparison, SNP-$h^2$ derived using LD Score Regression (LDSC) are also shown.

ǂ Estimated sample size based on the increase in mean $\chi^2$ statistic using multi-trait analysis of genome-wide association[38].

Abbreviations: EV, expressive vocabulary; ERV, expressive and receptive vocabulary; GWAS, genome-wide association study; $N_{ind}$ – Number of individuals; RV, receptive vocabulary

Table S6: Comparison of beta coefficients and corresponding standard errors for MTAG-derived analyses

| MTAG analysis | Max. FDR | Fixed effect meta-analysis | All SNPs | | | $P$<5x10$^{-6}$ SNPs | | |
|---|---|---|---|---|---|---|---|---|
| | | | N | $r_{\beta}$ | $r_{SE}$ | N | $r_{\beta}$ | $r_{SE}$ |
| Late-phase EV (stage I) | 0.42 | Late-phase EV excl. ALSPAC 38m | 7,355,069 | 0.84 | 0.97 | 37 | 0.99 | >0.99 |
| EV (stage II) | 0.41 | Late-phase EV excl. ALSPAC 38m | 7,343,861 | 0.84 | 0.97 | 37 | 0.99 | >0.99 |
| Late-phase ERV (stage II) | 0.38 | Late-phase EV excl. ALSPAC 38m | 7,355,069 | 0.77 | 0.97 | 37 | 0.99 | >0.99 |

For each multi-trait analysis of genome-wide association (MTAG) meta-analysis, beta coefficients and standard errors were compared with corresponding estimates from fixed-effect meta-analyses for late-phase expressive vocabulary (stage I, excluding ALSPAC expressive vocabulary at 38 months) across all shared SNP signals ($N_{SNPs}$=7,343,861-7,355,069) andacross a subset of highly-associated SNPs ($P$<5x10$^{-6}$, $N_{SNPs}$=37) using Pearson correlations. Meta-analysis SNP estimates were robust, despite high maximum false discovery rates.

Abbreviations: ALSPAC, Avon Longitudinal Study of Parents and Children; EV, expressive vocabulary; FDR, false discovery rate; m, months; max, maximum; MTAG, multi-trait analysis of genome-wide association; SNPs, single-nucleotide polymorphism; ERV, expressive/receptive vocabulary

Table S7: SNP-heritability of later-life outcomes included in High-Definition Likelihood analyses

| Trait | Reference panel | SNP-$h^2$(SE) | Z-score | *P* | $N_{ind}$ |
|---|---|---|---|---|---|
| Intelligence | HapMap3 | 0.17(5x$10^{-3}$) | 6.29 | <$10^{-10}$ | 279,930 |
| Educational attainment | HapMap3 | 0.10(2x$10^{-3}$) | 43.71 | <$10^{-10}$ | 766,345 |
| Infant head circumference | HapMap2 | 0.27(0.02) | 12.03 | <$10^{-10}$ | 10,768 |
| Childhood head circumference | HapMap3 | 0.26(0.04) | 6.29 | 3x$10^{-10}$ | 10,600 |
| Childhood aggression | HapMap3 | 0.03(3x$10^{-3}$) | 9.79 | <$10^{-10}$ | 151,741 |
| Childhood internalising symptoms | HapMap3 | 0.03(7x$10^{-3}$) | 4.50 | 8x$10^{-6}$ | 64,641 |
| ADHD | HapMap2 | 0.22(0.01) | 31.28 | <$10^{-10}$ | 53,293 |
| ASD | HapMap3 | 0.23(0.01) | 26.48 | <$10^{-10}$ | 46,350 |

SNP-heritability, corresponding standard error and *P*-value were estimated with HDL[47]. SNP-$h^2$ Z-scores were calculated by dividing SNP-$h^2$ by its standard error.

Abbreviations: ADHD, Attention-Deficit/Hyperactivity Disorder; ASD, Autism Spectrum Disorder; $N_{ind}$ – Number of individuals

Table S8: Power calculations for genetic overlap of early-life vocabulary with educational attainment

| Trait | $r_g$ value | | | | | | | | | |
|---|---|---|---|---|---|---|---|---|---|---|
| | 0.1 | 0.2 | 0.3 | 0.4 | 0.5 | 0.6 | 0.7 | 0.8 | 0.9 | 1.0 |
| Early-phase EV | 0.46 | 1 | 1 | 1 | 1 | 1 | 1 | 1 | 1 | 1 |
| Late-phase EV | 0.31 | 0.96 | 1 | 1 | 1 | 1 | 1 | 1 | 1 | 1 |
| Late-phase RV | 0.24 | 0.91 | 1 | 1 | 1 | 1 | 1 | 1 | 1 | 1 |

Statistical power to detect genetic overlap between single-trait vocabulary summary statistics and educational attainment was calculated online (https://eagenetics.shinyapps.io/power_website/) following Dudbridge et al[64]. The exact parameters used to derive power estimates are reported in the Supplemental Methods.

Abbreviations: EV, expressive vocabulary; RV, receptive vocabulary; $r_g$, genetic correlation

Table S9: Cholesky decomposition of early-life vocabulary size and later ADHD symptoms

| Genetic (co)variance | | | Residual (co)variance | | |
|---|---|---|---|---|---|
| Label | Path coefficient (SE) | *P* | Label | Path coefficient (SE) | *P* |
| a11 | 0.34(0.07) | $1.9x10^{-6}$ | e11 | 0.94(0.03) | $<1x10^{-10}$ |
| a21 | 0.20(0.10) | 0.04 | e21 | 0.50(0.04) | $<1x10^{-10}$ |
| a31 | 0.14(0.11) | 0.21 | e31 | 0.22(0.04) | $4.6x10^{-8}$ |
| a41 | -0.02(0.10). | 0.84 | e41 | 0.24(0.04) | $4.4x10^{-11}$ |
| a51 | 0.28(0.13) | 0.03 | e51 | -0.16(0.05) | 0.001 |
| a61 | 0.27(0.12) | 0.03 | e61 | -0.14(0.04) | 0.002 |
| a22 | 0.33(0.06) | $1.7x10^{-8}$ | e22 | 0.78(0.03) | $<1x10^{-10}$ |
| a32 | 0.28(0.09) | 0.002 | e32 | 0.32(0.04) | $<1x10^{-10}$ |
| a42 | 0.31(0.08) | $9.0x10^{-5}$ | e42 | 0.22(0.04) | $3.4x10^{-9}$ |
| a52 | -0.41(0.11) | $3.4x10^{-4}$ | e52 | 0.04(0.05) | 0.46 |
| a62 | -0.25(0.12) | 0.05 | e62 | -0.01(0.05) | 0.75 |
| a33 | 0.30(0.08) | $3.1x10^{-4}$ | e33 | 0.81(0.03) | $<1x10^{-10}$ |
| a43 | 0.14(0.11) | 0.20 | e43 | 0.46(0.03) | $<1x10^{-10}$ |
| a53 | 0.04(0.20) | 0.85 | e53 | -0.11(0.06) | 0.05 |
| a63 | -0.03(0.17) | 0.84 | e63 | -0.04(0.05) | 0.45 |
| a44 | 0.15(0.08) | 0.07 | e44 | 0.74(0.02) | 0 |
| a54 | 0.09(0.19) | 0.62 | e54 | 0.04(0.05) | 0.43 |
| a64 | -0.34(0.14) | 0.01 | e64 | 0.08(0.04) | 0.06 |
| a55 | $-7.1x10^{-5}$(NA) | NA | e55 | 0.84(0.05) | $<1x10^{-10}$ |
| a65 | $-8.7x10^{-6}$(0.31) | 1.00 | e65 | 0.24(0.07) | $3.0x10^{-4}$ |
| a66 | $-7.8x10^{-5}$(0.19) | 1.00 | e66 | 0.82(0.04) | $<1x10^{-10}$ |

Genetic-relationship matrix structural equation modelling (GRM-SEM) of early-life vocabulary scores (15, 24 and 38 months of age) and ADHD symptom scores (teacher-report at 8 years and mother-report at 13 years) based on all available observations for children across development (N≤6,524). Path coefficients originating from genetic factors are labelled with ‘a’, whereas path coefficients originating from residual factors are labelled with ‘e’. The first number indicates the measure onto which the factor loads, while the second number indicates the respective factor. Thus, a21 refers to the path from factor A1, capturing genetic variance in early-phase expressive vocabulary to late-phase expressive vocabulary (Figure 4a). Individual-level data was retrieved from the Avon Longitudinal Study of Parents and Children.

### Supplemental figures

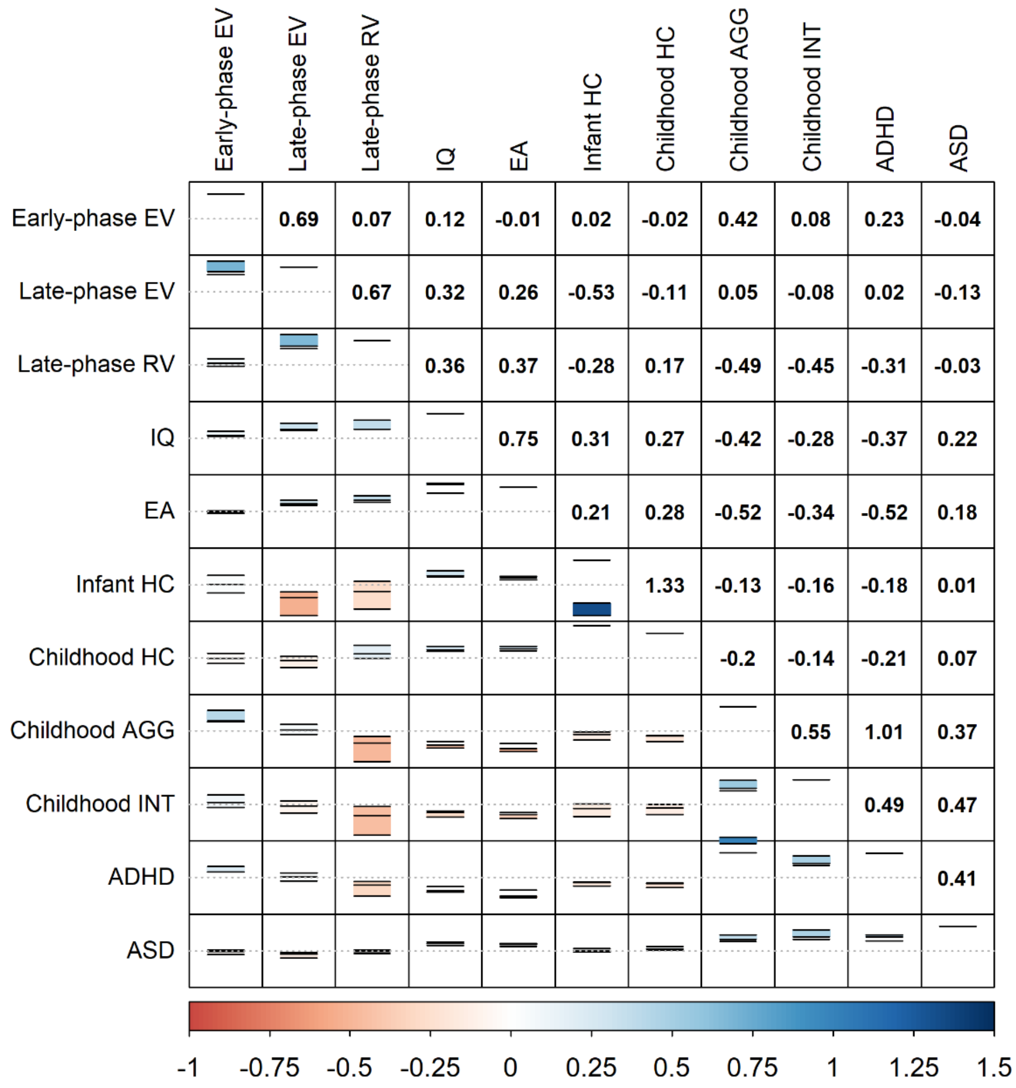

**Figure S1: Genetic correlations among traits included in High-Definition Likelihood analyses**

Genetic correlations ($r_g$) were estimated using genome-wide summary statistics and High-Definition Likelihood[47]. The lower triangle represents $r_g$ estimates and corresponding standard errors, with the dotted line representing an estimate of zero. The upper triangle represents $r_g$ estimates in number format.

Abbreviatons: ADHD, Attention-Deficit/Hyperactivity Disorder; AGG, aggression; ASD, Autism Spectrum Disorder; EA, educational attainment; EV, expressive vocabulary; HC, head circumference; INT, internalising symptoms; IQ, general intelligence; RV, receptive vocabulary

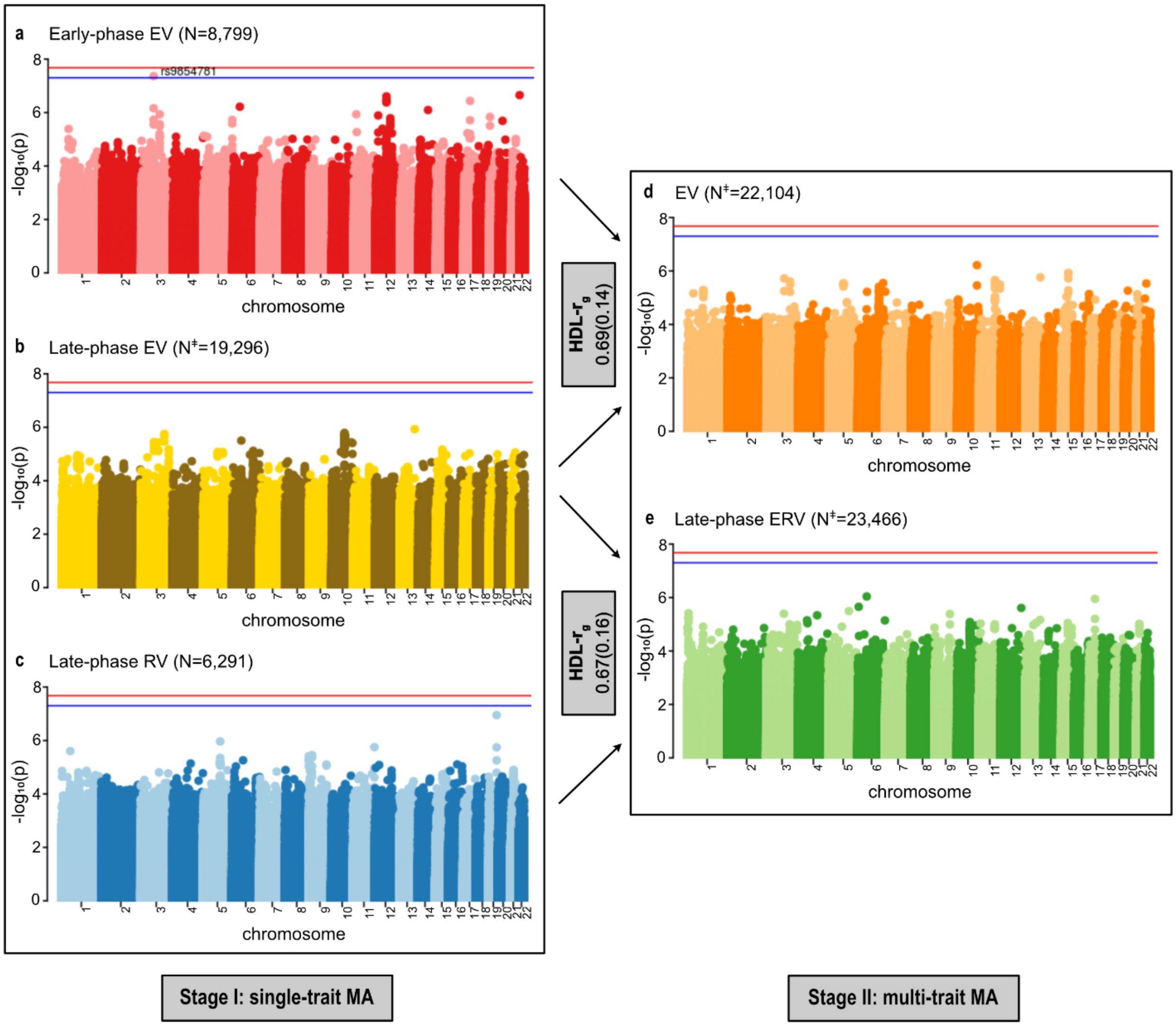

**Figure S2: Genome-wide association results for single-trait and multi-trait vocabulary meta-analyses**

Genome-wide association with **(a)** early-phase expressive vocabulary (N=8,799); **(b)** late-phase expressive vocabulary (N‡=19,296) and **(c)** receptive vocabulary (N=6,291), as estimated using single-trait meta-analyses as part of stage I. Multi-trait genome-wide association results (stage II) estimated using MTAG for **(d)** expressive vocabulary (N‡=22,104), representing early- and late-phase expressive vocabulary (stage I) and **(e)** late-phase expressive and receptive vocabulary (N‡=23,466), representing late-phase expressive and receptive vocabulary (stage I). No associations passed the genome-wide significance threshold adjusted for the number of independent traits studied of $2.10 \times 10^{-8}$ (red line). The blue line represents the unadjusted genome-wide significance threshold of $5 \times 10^{-8}$, variants passing this threshold are labelled in black. Genomic positions are shown according to NCBI Build 37. Genetic correlations between single-trait vocabulary summary statistics were derived using high-definition likelihood.

‡ Estimated sample size based on the increase in mean $\chi^2$ statistic using multi-trait analysis of genome-wide association.

Abbreviations: EV, expressive vocabulary; ERV, expressive and receptive vocabulary; HDL, High-Definition Likelihood; MA, meta-analyses; N, sample size; $r_g$, genetic correlation; RV, receptive vocabulary

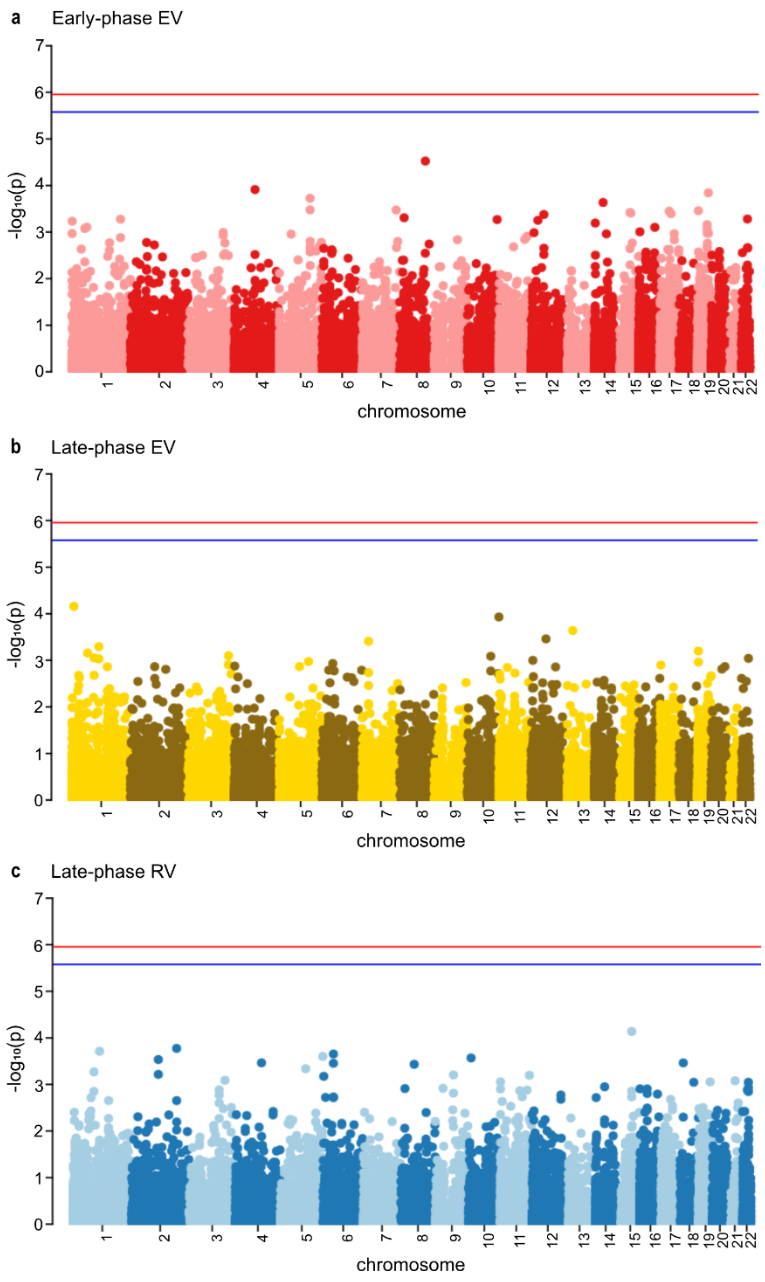

**Figure S3: Gene-based GWASs**

Gene-based genome-wide association analyses for single-trait vocabulary summary statistics of **(a)** early-phase expressive vocabulary, **(b)** late-phase expressive vocabulary, and **(c)** late-phase receptive vocabulary. No associations passed the genome-wide significance threshold adjusted for the number of genes and independent traits studied of $1.11\text{x}10^{-6}$ (red line). The blue line represents the unadjusted genome-wide significance threshold of $2.64\text{x}10^{-6}$. Genomic positions are shown according to NCBI Build 37.

Abbreviations: EV, expressive vocabulary; RV, receptive vocabulary

### Supplemental References

1. Clark EV. *First Language Acquisition*. Cambridge University Press; 2016.

2. Fenson L, Dale PS, Reznick JS, Bates E, Thal DJ, Pethick SJ. Variability in early communicative development. *Monogr Soc Res Child Dev*. 1994;59(5):1-173; discussion 174-85.

3. Hoff E. *Language Development*. Cengage Learning; 2013.

4. Houston-Price C, Mather E, Sakkalou E. Discrepancy between parental reports of infants' receptive vocabulary and infants' behaviour in a preferential looking task. *J Child Lang*. 2007;34(4):701-724. doi:10.1017/s0305000907008124

5. Verhoef E, Shapland CY, Fisher SE, Dale PS, St Pourcain B. The developmental genetic architecture of vocabulary skills during the first three years of life: Capturing emerging associations with later-life reading and cognition. *PLOS Genetics*. 2021;17(2):e1009144. doi:10.1371/journal.pgen.1009144

6. Boyd A, Golding J, Macleod J, et al. Cohort Profile: the 'children of the 90s'--the index offspring of the Avon Longitudinal Study of Parents and Children. *Int J Epidemiol*. 2013;42(1):111-127. doi:10.1093/ije/dys064

7. Fraser A, Macdonald-Wallis C, Tilling K, et al. Cohort Profile: the Avon Longitudinal Study of Parents and Children: ALSPAC mothers cohort. *Int J Epidemiol*. 2013;42(1):97-110. doi:10.1093/ije/dys066

8. Vuillermin P, Saffery R, Allen KJ, et al. Cohort Profile: The Barwon Infant Study. *Int J Epidemiol*. 2015;44(4):1148-1160. doi:10.1093/ije/dyv026

9. Bjarnadóttir E, Stokholm J, Chawes B, et al. Determinants of neurodevelopment in early childhood - results from the Copenhagen prospective studies on asthma in childhood (COPSAC2010 ) mother-child cohort. *Acta Paediatr*. 2019;108(9):1632-1641. doi:10.1111/apa.14753

10. Ahluwalia TS, Eliasen AU, Sevelsted A, et al. FUT2-ABO epistasis increases the risk of early childhood asthma and Streptococcus pneumoniae respiratory illnesses. *Nat Commun*. 2020;11(1):6398. doi:10.1038/s41467-020-19814-6

11. Reilly S, Cook F, Bavin EL, et al. Cohort Profile: The Early Language in Victoria Study (ELVS). *International Journal of Epidemiology*. 2018;47(1):11-20. doi:10.1093/ije/dyx079

12. Kooijman MN, Kruithof CJ, van Duijn CM, et al. The Generation R Study: design and cohort update 2017. *European Journal of Epidemiology*. 2016;31(12):1243-1264. doi:10.1007/s10654-016-0224-9

13. Medina-Gomez C, Felix JF, Estrada K, et al. Challenges in conducting genome-wide association studies in highly admixed multi-ethnic populations: the Generation R Study. *European Journal of Epidemiology*. 2015;30(4):317-330. doi:10.1007/s10654-015-9998-4

14. Soloff C, Lawrence D, Johnstone R. *The Longitudinal Study of Australian Children: Sample Design: LSAC Technical Paper Number 1.*; 2005:30. https://api.research-repository.uwa.edu.au/ws/portalfiles/portal/73664759/tp1.pdf

15. Clifford SA, Davies S, Wake M, Child Health CheckPoint Team. Child Health CheckPoint: cohort summary and methodology of a physical health and biospecimen module for the Longitudinal Study of Australian Children. *BMJ Open*. 2019;9(Suppl 3). doi:10.1136/bmjopen-2017-020261

16. Newnham JP, Evans SF, Michael CA, Stanley FJ, Landau LI. Effects of frequent ultrasound during pregnancy: a randomised controlled trial. *The Lancet*. 1993;342(8876):887-891. doi:10.1016/0140-6736(93)91944-H

17. Straker L, Mountain J, Jacques A, et al. Cohort Profile: The Western Australian Pregnancy Cohort (Raine) Study–Generation 2. *International Journal of Epidemiology*. 2017;46(5):dyw308. doi:10.1093/ije/dyw308

18. Rimfeld K, Malanchini M, Spargo T, et al. Twins Early Development Study: A Genetically Sensitive Investigation into Behavioral and Cognitive Development from Infancy to Emerging Adulthood. *Twin Research and Human Genetics*. 2019;22(6):508-513. doi:10.1017/thg.2019.56

19. Fenson L, Dale P, Reznick JS, et al. *User's Guide and Technical Manual for the MacArthur Communicative Development Inventories*. Singular Publishing; 1993.

20. Rescorla Leslie. The Language Development Survey. *Journal of Speech and Hearing Disorders*. 1989;54(4):587-599. doi:10.1044/jshd.5404.587

21. Bleses D, Vach W, Slott M, et al. The Danish Communicative Developmental Inventories: validity and main developmental trends. *J Child Lang*. 2008;35(3):651-669. doi:10.1017/S0305000907008574

22. Zink I, Lejaegere M. N-CDI's: korte vormen, Aanpassing en hernormering van de MacArthur Short Form Vocabulary Checklist van Fenson et al.

23. Dale PS, Goodman JC. Commonality and Individual Differences in Vocabulary Growth. In: *Beyond Nature-Nurture: Essays in Honor of Elizabeth Bates*. Lawrence Erlbaum Associates Publishers; 2005:41-78.

24. Bleses D, Vach W, Slott M, et al. Early vocabulary development in Danish and other languages: a CDI-based comparison. *J Child Lang*. 2008;35(3):619-650. doi:10.1017/S0305000908008714

25. Fenson L, Pethick S, Renda C, Cox JL, Dale PS, Reznick JS. Short-form versions of the MacArthur Communicative Development Inventories. *Applied Psycholinguistics*. 2000;21(01):95-116. doi:null

26. Dale PS, Dionne G, Eley TC, Plomin R. Lexical and grammatical development: a behavioural genetic perspective. *Journal of Child Language*. 2000;27(03):619-642. doi:null

27. Fenson L, Marchman VA. *MacArthur-Bates Communicative Development Inventories: User's Guide and Technical Manual*. Paul H. Brookes Publishing Company; 2007.

28. Rescorla L, Ratner NB, Jusczyk P, Jusczyk AM. Concurrent validity of the language development survey: associations with the MacArthur-Bates communicative development inventories: words and sentences. *Am J Speech Lang Pathol*. 2005;14(2):156-163. doi:10.1044/1058-0360(2005/016)

29. Marchini J, Howie B, Myers S, McVean G, Donnelly P. A new multipoint method for genome-wide association studies by imputation of genotypes. *Nat Genet*. 2007;39(7):906-913. doi:10.1038/ng2088

30. Aulchenko YS, Struchalin MV, van Duijn CM. ProbABEL package for genome-wide association analysis of imputed data. *BMC Bioinformatics*. 2010;11:134. doi:10.1186/1471-2105-11-134

31. Zhou X, Stephens M. Genome-wide efficient mixed-model analysis for association studies. *Nat Genet*. 2012;44(7):821-824. doi:10.1038/ng.2310

32. Chang CC, Chow CC, Tellier LC, Vattikuti S, Purcell SM, Lee JJ. Second-generation PLINK: rising to the challenge of larger and richer datasets. *Gigascience*. 2015;4(1). doi:10.1186/s13742-015-0047-8

33. Tucker G, Loh PR, MacLeod IM, et al. Two-Variance-Component Model Improves Genetic Prediction in Family Datasets. *Am J Hum Genet*. 2015;97(5):677-690. doi:10.1016/j.ajhg.2015.10.002

34. Zaitlen N, Kraft P, Patterson N, et al. Using Extended Genealogy to Estimate Components of Heritability for 23 Quantitative and Dichotomous Traits. *PLoS Genet*. 2013;9(5). doi:10.1371/journal.pgen.1003520

35. Winkler TW, Day FR, Croteau-Chonka DC, et al. Quality control and conduct of genome-wide association meta-analyses. *Nat Protoc*. 2014;9(5):1192-1212. doi:10.1038/nprot.2014.071

36. Devlin B, Roeder K. Genomic control for association studies. *Biometrics*. 1999;55(4):997-1004.

37. Willer CJ, Li Y, Abecasis GR. METAL: fast and efficient meta-analysis of genomewide association scans. *Bioinformatics*. 2010;26(17):2190-2191. doi:10.1093/bioinformatics/btq340

38. Turley P, Walters RK, Maghzian O, et al. Multi-trait analysis of genome-wide association summary statistics using MTAG. *Nature Genetics*. 2018;50(2):229-237. doi:10.1038/s41588-017-0009-4

39. Purcell S, Cherny SS, Sham PC. Genetic Power Calculator: design of linkage and association genetic mapping studies of complex traits. *Bioinformatics*. 2003;19(1):149-150. doi:10.1093/bioinformatics/19.1.149

40. Leeuw CA de, Mooij JM, Heskes T, Posthuma D. MAGMA: Generalized Gene-Set Analysis of GWAS Data. *PLOS Computational Biology*. 2015;11(4):e1004219. doi:10.1371/journal.pcbi.1004219

41. Watanabe K, Taskesen E, Bochoven A van, Posthuma D. Functional mapping and annotation of genetic associations with FUMA. *Nature Communications*. 2017;8(1):1826. doi:10.1038/s41467-017-01261-5

42. Nyholt DR. A Simple Correction for Multiple Testing for Single-Nucleotide Polymorphisms in Linkage Disequilibrium with Each Other. *Am J Hum Genet*. 2004;74(4):765-769.

43. Subramanian A, Tamayo P, Mootha VK, et al. Gene set enrichment analysis: a knowledge-based approach for interpreting genome-wide expression profiles. *Proc Natl Acad Sci USA*. 2005;102(43):15545-15550. doi:10.1073/pnas.0506580102

44. Mooney MA, Wilmot B. Gene Set Analysis: A Step-By-Step Guide. *Am J Med Genet B Neuropsychiatr Genet*. 2015;168(7):517-527. doi:10.1002/ajmg.b.32328

45. Aguet F, Barbeira AN, Bonazzola R, et al. The GTEx Consortium atlas of genetic regulatory effects across human tissues. *bioRxiv*. Published online October 3, 2019:787903. doi:10.1101/787903

46. Li M, Santpere G, Imamura Kawasawa Y, et al. Integrative functional genomic analysis of human brain development and neuropsychiatric risks. *Science*. 2018;362(6420). doi:10.1126/science.aat7615

47. Ning Z, Pawitan Y, Shen X. High-definition likelihood inference of genetic correlations across human complex traits. *Nat Genet*. 2020;52(8):859-864. doi:10.1038/s41588-020-0653-y

48. Savage JE, Jansen PR, Stringer S, et al. Genome-wide association meta-analysis in 269,867 individuals identifies new genetic and functional links to intelligence. *Nature Genetics*. 2018;50(7):912-919. doi:10.1038/s41588-018-0152-6

49. Lee JJ, Wedow R, Okbay A, et al. Gene discovery and polygenic prediction from a genome-wide association study of educational attainment in 1.1 million individuals. *Nature Genetics*. 2018;50(8):1112-1121. doi:10.1038/s41588-018-0147-3

50. Taal HR, St Pourcain B, Thiering E, et al. Common variants at 12q15 and 12q24 are associated with infant head circumference. *Nat Genet*. 2012;44(5):532-538. doi:10.1038/ng.2238

51. Haworth S, Shapland CY, Hayward C, et al. Low-frequency variation in TP53 has large effects on head circumference and intracranial volume. *Nature Communications*. 2019;10(1):357. doi:10.1038/s41467-018-07863-x

52. Ip HF, van der Laan CM, Krapohl EML, et al. Genetic association study of childhood aggression across raters, instruments, and age. *Transl Psychiatry*. 2021;11(1):1-9. doi:10.1038/s41398-021-01480-x

53. Jami ES, Hammerschlag AR, Ip HF, et al. Genome-wide association meta-analysis of childhood and adolescent internalising symptoms. Published online July 31, 2021:2020.09.11.20175026. Accessed August 12, 2021. https://www.medrxiv.org/content/10.1101/2020.09.11.20175026v2

54. Demontis D, Walters RK, Martin J, et al. Discovery of the first genome-wide significant risk loci for attention deficit/hyperactivity disorder. *Nature Genetics*. Published online November 26, 2018:1. doi:10.1038/s41588-018-0269-7

55. *International Statistical Classification of Diseases and Related Health Problems*. Vol 10th revision. World Health Organization; 2010.

56. Mors O, Perto GP, Mortensen PB. The Danish Psychiatric Central Research Register. *Scand J Public Health*. 2011;39(7 Suppl):54-57. doi:10.1177/1403494810395825

57. American Psychiatric Association. *Diagnostic and Statistical Manual of Mental Disorders*. Vol fifth. American Psychiatric Association; 2013.

58. Grove J, Ripke S, Als TD, et al. Identification of common genetic risk variants for autism spectrum disorder. *Nature Genetics*. 2019;51(3):431. doi:10.1038/s41588-019-0344-8

59. Pedersen CB, Bybjerg-Grauholm J, Pedersen MG, et al. The iPSYCH2012 case–cohort sample: new directions for unravelling genetic and environmental architectures of severe mental disorders. *Molecular Psychiatry*. 2018;23(1):6-14. doi:10.1038/mp.2017.196

60. Cross-Disorder Group of the Psychiatric Genomics Consortium. Genetic relationship between five psychiatric disorders estimated from genome-wide SNPs. *Nat Genet*. 2013;45(9):984-994. doi:10.1038/ng.2711

61. Lord C, Rutter M, Le Couteur A. Autism Diagnostic Interview-Revised: a revised version of a diagnostic interview for caregivers of individuals with possible pervasive developmental disorders. *J Autism Dev Disord*. 1994;24(5):659-685.

62. Lord C, Rutter M, Goode S, et al. Autism diagnostic observation schedule: a standardized observation of communicative and social behavior. *J Autism Dev Disord*. 1989;19(2):185-212. doi:10.1007/BF02211841

63. Berument SK, Rutter M, Lord C, Pickles A, Bailey A. Autism screening questionnaire: diagnostic validity. *Br J Psychiatry*. 1999;175:444-451. doi:10.1192/bjp.175.5.444

64. Dudbridge F. Power and Predictive Accuracy of Polygenic Risk Scores. *PLOS Genet*. 2013;9(3):e1003348. doi:10.1371/journal.pgen.1003348

65. Reilly S, Bavin EL, Bretherton L, et al. The Early Language in Victoria Study (ELVS): A prospective, longitudinal study of communication skills and expressive vocabulary development at 8, 12 and 24 months. *International Journal of Speech-Language Pathology*. 2009;11(5):344-357. doi:10.1080/17549500903147560

66. Marees AT, Kluiver H de, Stringer S, et al. A tutorial on conducting genome-wide association studies: Quality control and statistical analysis. *International Journal of Methods in Psychiatric Research*. 2018;27(2):e1608. doi:10.1002/mpr.1608

67. McCarthy S, Das S, Kretzschmar W, et al. A reference panel of 64,976 haplotypes for genotype imputation. *Nat Genet*. 2016;48(10):1279-1283. doi:10.1038/ng.3643

68. Ge T, Chen CY, Ni Y, Feng YCA, Smoller JW. Polygenic prediction via Bayesian regression and continuous shrinkage priors. *Nat Commun*. 2019;10(1):1776. doi:10.1038/s41467-019-09718-5

69. St Pourcain B, Eaves LJ, Ring SM, et al. Developmental changes within the genetic architecture of social communication behaviour: A multivariate study of genetic variance in unrelated individuals. *Biological Psychiatry*. 2017;83:598-606. doi:10.1016/j.biopsych.2017.09.020

70. Grotzinger AD, Rhemtulla M, de Vlaming R, et al. Genomic structural equation modelling provides insights into the multivariate genetic architecture of complex traits. *Nature Human Behaviour*. 2019;3(5):513-525. doi:10.1038/s41562-019-0566-x

71. Middeldorp CM, Hammerschlag AR, Ouwens KG, et al. A Genome-Wide Association Meta-Analysis of Attention-Deficit/Hyperactivity Disorder Symptoms in Population-Based Pediatric Cohorts. *J Am Acad Child Adolesc Psychiatry*. 2016;55(10):896-905.e6. doi:10.1016/j.jaac.2016.05.025

72. Goodman R, Ford T, Simmons H, Gatward R, Meltzer H. Using the Strengths and Difficulties Questionnaire (SDQ) to screen for child psychiatric disorders in a community sample. *Int Rev Psychiatry*. 2003;15(1-2):166-172. doi:10.1080/0954026021000046128

73. Yang J, Benyamin B, McEvoy BP, et al. Common SNPs explain a large proportion of the heritability for human height. *Nat Genet*. 2010;42(7):565-569. doi:10.1038/ng.608

74. Lee SH, Yang J, Goddard ME, Visscher PM, Wray NR. Estimation of pleiotropy between complex diseases using single-nucleotide polymorphism-derived genomic relationships and restricted maximum likelihood. *Bioinformatics*. 2012;28(19):2540-2542. doi:10.1093/bioinformatics/bts474

75. Yang J, Lee SH, Goddard ME, Visscher PM. GCTA: a tool for genome-wide complex trait analysis. *Am J Hum Genet*. 2011;88(1):76-82. doi:10.1016/j.ajhg.2010.11.011

76. Schlag F, Allegrini AG, Buitelaar J, et al. Polygenic risk for mental disorder reveals distinct association profiles across social behaviour in the general population. *Mol Psychiatry*. Published online February 28, 2022:1-11. doi:10.1038/s41380-021-01419-0

77. Neale M, Boker S, Xie G, Meas HHM. *Mx: Statistical Modeling*. 7th ed. Department of Psychiatry, Medical College of Virginia; 2006.

78. Falconer DS, Mackay TFC. *Quantitative Genetics*. Vol fourth. Pearson; 1996.

79. Loh PR, Danecek P, Palamara PF, et al. Reference-based phasing using the Haplotype Reference Consortium panel. *Nat Genet*. 2016;48(11):1443-1448. doi:10.1038/ng.3679

80. Durbin R. Efficient haplotype matching and storage using the positional Burrows-Wheeler transform (PBWT). *Bioinformatics*. 2014;30(9):1266-1272. doi:10.1093/bioinformatics/btu014

81. Das S, Forer L, Schönherr S, et al. Next-generation genotype imputation service and methods. *Nat Genet*. 2016;48(10):1284-1287. doi:10.1038/ng.3656

82. Bulik-Sullivan BK, Loh PR, Finucane HK, et al. LD Score regression distinguishes confounding from polygenicity in genome-wide association studies. *Nature genetics*. 2015;47(3):291-295. doi:10.1038/ng.3211
